# Saxiphilin functions as a ‘toxin sponge’ protein that counteracts the effects of saxitoxin poisoning

**DOI:** 10.1101/2025.11.20.689596

**Authors:** Samantha A. Nixon, Sandra Zakrzewska, Seil Jang, Keli Huang, Anissa Bara, Zhou Chen, Daynen R. Goss, Elizabeth R. Park, J. Du Bois, Daniel L. Minor

## Abstract

Saxitoxin (STX) is among the most potent toxins known, is classified as a chemical weapon, and is the archetype of the paralytic shellfish toxin (PST) family produced by marine and freshwater harmful algal blooms (HAB)^1–3^. STX causes paralysis and death through inhibition of voltage-gated sodium channels (Na_V_s), has no antidote, and poses a public health and commercial fishing threat due to its accumulation in seafood and increasing HAB occurrences^1,4^. Although STX is lethal to varied vertebrates^5–7^, including humans^4,8,9^, certain frogs resist STX poisoning^6,7^. This phenotype is thought to depend on soluble ‘toxin sponge’ high-affinity STX binding proteins (saxiphilins, Sxphs)^6,7,10,11^ that provide resistance through a competition mechanism different from classic ‘target site’ ion channel mutation mechanisms^12–14^. Here, we test this idea directly and show that a single dose of American bullfrog (*Rana catesbeiana*) Sxph (*Rc*Sxph) is sufficient to counteract STX neurotoxicity and lethality in mice using paradigms where STX and Sxph are administered together or sequentially in either order. Importantly, this function requires the *Rc*Sxph high-affinity STX binding site. Our findings provide *in vivo* validation that Sxphs are ‘toxin sponges’ that protect against STX poisoning and highlight the potential to harness this protein class as antidotes for PSTs and other toxins.

## Introduction

Saxitoxin (STX) is one of the most potent neurotoxins on the planet^1^, and is the archetype of a group of over 50 related congeners that can accumulate in shellfish and hence, are known as paralytic shellfish toxins (PSTs)^1–3^. This small (∼300 Da), highly water-soluble bis-guanidinium toxin is produced by cyanobacteria and dinoflagellates that cause Harmful Algal Blooms (HABs) in marine and freshwater, collectively known as ‘red tides’^1,3^. The neurotoxicity of STX arises from its potent capacity to block voltage-gated sodium channels (Na_V_s) that prevents action potential propagation in nerves and muscles causing paralysis and death^1,4^. Its toxicity is so severe that it is the only marine toxin to be classified as a chemical weapon^1,4^. Because of its threat to commercial fisheries and public health and the increased frequency of HABs^15–17^, worldwide programs to monitor PSTs to ensure food security are becoming ever more important^18,19^. Consumption of PST-contaminated shellfish can cause paralytic shellfish poisoning (PSP) in as little as 30 minutes leading to nausea, vomiting, numbness, paresthesia, dysphagia, ataxia, hypertension, and paralysis^9,20,21^. The rapid paralysis induced by PSP can progress to respiratory failure and death, with some fatalities reported within 3–4 hours post-ingestion^8,9^. Although there has been a long standing interest in developing treatments for PSP ranging from antibody-based approaches^22–24^ to neuron-derived nanoparticles^25^, to date, no antidote exists for STX or any other PST, and treatment relies upon supportive care and respiratory intervention^9,21^. Hence, there remains a need to develop means to counter the neurotoxic and lethal effects of PSTs.

Saxiphilins (Sxphs) are a family of soluble, ∼90-120 kDa, high affinity STX binding proteins found in anurans (frogs and toads)^11,26–28^. These ‘toxin sponge’ proteins^6^ are thought to be responsible for the ability of certain frog species to resist the lethal effects of STX poisoning^6,7,10,11^, even surviving injections of up to 20× the median lethal dose (LD_50_) for mice^6^. Sxphs have a single high affinity STX binding site (single digit nM Kd)^11,26,27,29,30^ with an affinity for STX that matches Na_V_s^5,30^. Electrophysiological experiments have shown that binding of STX and STX congeners to this site endows Sxphs with the ability to rescue Na_V_s from block by these toxins^26,30^. The capacity of Sxphs to act as ‘toxin sponges’ that effectively compete with Na_V_s for toxin binding not only suggests a biological role for Sxphs in frog STX resistance mechanisms^6,11^, but also raises the possibility that Sxph could be used as a countermeasure against STX poisoning.

Here, using mouse models of STX poisoning, we test whether the Sxph *in vitro* ‘toxin sponge’ functionality translates into an effective anti-STX agent *in vivo*. Using *in vitro* assays, we find that the prototypical 91 kDa Sxph from the American bullfrog (*Rana catesbeiana*) (*Rc*Sxph)^11,26,27^ rescues human Na_V_ channels from STX block, and that the high-affinity of Sxph:STX interaction is stably maintained across a wide temperature range that includes typical mammalian body temperatures. Strikingly, a single dose of *Rc*Sxph is efficacious for *in vivo* protection from the lethal effects of STX poisoning in mice in three separate poisoning paradigms in which STX and Sxph are administered together or sequentially in either order. Further, we observe that *Rc*Sxph administration not only prevents STX-induced death but also ameliorates the frequency and severity of STX poisoning symptoms, and has no adverse effects on its own. Prevention of the neurotoxic and lethal effects of STX intoxication is entirely dependent on the capacity of *Rc*Sxph to bind STX with high affinity, matching the outcomes of *in vitro* studies^26^. Together, our data provide the first direct evidence that Sxph alone is sufficient to protect an organism from STX poisoning and validate the idea that these proteins act as ‘toxin sponges’ capable of mitigating the physiological effects of toxins^6^. These findings, together with the capacity of Sxphs to bind different classes of STX congeners^30,31^, highlight the possibility of harnessing this class of proteins as a platform for creating antitoxins against PSP.

## Results

### *Rc*Sxph protects multiple human sodium channel isoforms from STX block

Functional characterization has shown that *Rc*Sxph can rescue amphibian Na_V_1.4 channels from block by STX^6,26^ and multiple STX congeners^30^. Similar studies demonstrated that a related Sxph from the High Himalaya frog (*Nanorana parkeri*), *Np*Sxph, exhibits the same behavior against mammalian Na_V_1.4^30^, highlighting the potential versatility these proteins as countermeasures against bis-guanidinium toxins. To assess whether *Rc*Sxph is also effective for rescuing mammalian Na_V_ channels, we used planar patch clamp electrophysiology. We compared the capacities of *Rc*Sxph and *Rc*Sxph E540A (henceforth E540A), a mutant in which STX affinity is decreased by ∼2000-fold^26^, to counteract STX block of three exemplar human Na_V_s representing skeletal muscle (*Hs*Na_V_1.4), cardiac (*Hs*Na_V_1.5), and peripheral nervous system (*Hs*Na_V_1.7) isoforms. We first measured the half-maximal inhibitory concentration (IC_50_) for STX against stable cell lines expressing each of these target channels (Fig. S1, Table S1). In agreement with previous studies^5,30,32^, STX was most potent against *Hs*Na_V_1.4 (IC_50_ = 3.5 ± 2.5 nM), and displayed ∼80-fold and ∼200-fold lower activity against *Hs*Na_V_1.5 and *Hs*Na_V_1.7, respectively (IC_50_= 284.6 ± 50.6 nM and 637.9 ± 189.5 nM, respectively). We then measured the capacity of a 3:1 [*Rc*Sxph]:[STX] molar ratio to counteract STX applied at concentrations sufficient to block ∼80% of the current for each channel isoform (Table S1). Sequential application of saline, STX, *Rc*Sxph:STX, STX, and *Rc*Sxph showed that *Rc*Sxph neutralized the effects of STX on all three human Na_V_ isoforms tested (Figs. 1A-B, D-E, and G-H). Using a ration of 3:1 [*Rc*Sxph]:[STX] resulted in near complete recovery of *Hs*Na_V_1.4 (mean current recovery 83.9 ± 7.3%) and complete restoration of *Hs*Na_V_1.5 and *Hs*Na_V_1.7 currents (109.6 ± 4.4%, and 96.9 ± 9.2% respectively) (Fig. 1J). By contrast, E540A had no effect on STX block of *Hs*Na_V_1.4 or *Hs*Na_V_1.5 (Fig. 1A and C, D and F), and afforded marginal restoration of *Hs*Na_V_1.7 currents (Fig. 1G and I) (mean current recovery −0.8 ± 0.8%, −7.0 ± 4.8%, 15.7 ± 13.8%, respectively) (Fig. 1J), aligning with its dramatically reduced STX affinity ^26^. These results establish that *Rc*Sxph shares the ability with *Np*Sxph to protect human Na_V_1.4, in line with their similar affinities for STX^26^, and can rescue multiple human Na_V_ isoforms from STX block, regardless of whether the channel has a high (Na_V_1.4) or low (Na_V_1.5 and Na_V_1.7) affinity for the toxin. Although E540A was generally ineffective at sequestering STX, we did observe a weak restorative effect (∼15%) of E540A on *Hs*Na_V_1.7 blocked with 3 µM STX (Fig. 1J). This result aligns with the estimated 16.4% toxin binding based on the reported E540A Kd (15.3 ± 4.1 µM)^26^ and the high concentration of both components used in this experiment relative to the Kd. Regardless, the overall ineffectiveness of E540A to rescue STX block of all three Na_V_s underscores that STX neutralization relies on high affinity interaction of STX with its binding site on Sxph^6,26,30^. Together, these data establish that Sxphs show broad capability for rescuing a range of Na_V_ isoforms from STX block and hence, should be able to suppress STX effects that arise from interaction with Na_V_s in different types of excitable cells.

**Figure 1:**
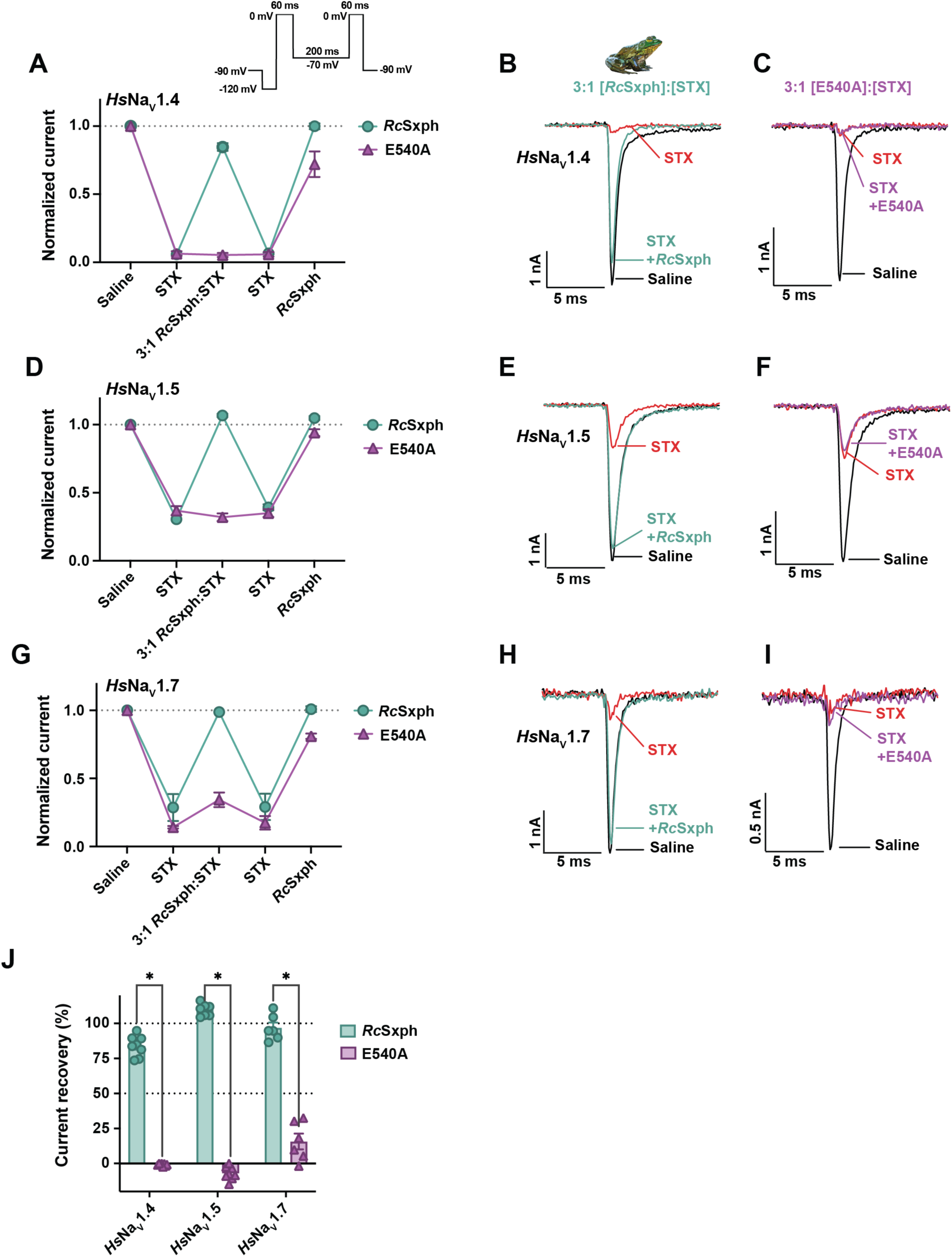
*Rc*Sxph protects multiple human Na_V_ channels from STX block *in vitro*. Sodium currents from cells stably expressing human Na_V_ isoforms *Hs*Na_V_1.4, *Hs*Na_V_1.5, and *Hs*Na_V_1.7 were recorded using automated planar whole-cell patch clamp electrophysiology. The capacity of *Rc*Sxph to protect channels from STX inhibition *in vitro* was assessed by comparing currents recorded under STX block (100 nM for *Hs*Na_V_1.4, 1.1 µM for *Hs*Na_V_1.5 and 3 µM for *Hs*Na_V_1.7) against currents exposed to equivalent concentration STX preincubated with a 3:1 molar ratio of Sxph:STX. The high-affinity wild-type *Rc*Sxph (green, circles) was compared against a low-affinity *Rc*Sxph mutant, E540A (purple, triangles), as a control. **A, D, G,** Baseline sodium currents were recorded in saline, and then cells were sequentially exposed to STX, followed by 3:1 [Sxph]:[STX], STX, and finally Sxph alone. Each point represents the mean peak sodium current recorded, with standard deviation (SD) for **A,** *Hs*Na_V_1.4 (*Rc*Sxph, *n* = 8; E540A, *n* = 6), **D,** *Hs*Na_V_1.5 (*Rc*Sxph, *n* = 7; E540A, *n* = 7), and **G,** *Hs*Na_V_1.7 (*Rc*Sxph, *n* = 6; E540A, *n* = 8). **B–C, E–F, H–I,** Corresponding exemplar whole-cell patch clamp recordings with baseline currents recorded in saline (black), in the presence of STX (red) and in the presence of either 3:1 [RcSxph]:[STX] (green) or 3:1 [E540A]:[STX] (purple), **B–C**, *Hs*Na_V_1.4, **E–F**, *Hs*Na_V_1.5, **H–I**, *Hs*Na_V_1.7. **J,** Current recovery (%) for 3:1 [*Rc*Sxph]:[STX] (green, circles) and 3:1 [E540A]:[STX] (purple, triangles). Error bars are standard error of the mean (SEM) and asterisks represent *p* < 0.0001 calculated by unpaired t-tests with Welch correction.

### *Rc*Sxph high-affinity binding to STX is thermally resilient

Studies of STX binding to *Rc*Sxph at ambient temperature (23 ± 2°C)^26,30^ or lower (0–25°C)^29^ show that STX affinity is not strongly perturbed over this temperature range. Before embarking on experiments to test the effectiveness of *Rc*Sxph as an STX countermeasure in mice where the body temperature is ∼36°C^33^, we wanted to examine the temperature dependence of STX binding at temperatures >25°C to understand if the elevated temperature would compromise interaction with the toxin. We used isothermal titration calorimetry (ITC) to measure STX binding to *Rc*Sxph at 35°C and 50°C and compared the results with our previous studies at 25°C^26^. The data show that, in line with the high stability of *Rc*Sxph, which has an apparent melting temperature (Tm) of ∼55°C in thermofluor (TF) assays^26^, the high affinity *Rc*Sxph:STX interaction is robustly maintained across the 25–50°C range (Table 1, Fig. S2). There are only minor changes in the measured thermodynamic parameters and affinity (Kd = 1.2 ± 0.8, 2.5 ± 1.4, and 7.5 ± 2.5 nM for 25°C, 35°C, and 50°C, respectively), yielding negligible binding energy differences between 25°C and 50°C (ΔΔG = 0.3 kcal mol^-1^, Table S2). The thermal resilience of *Rc*Sxph and high-affinity STX binding combined with its ability to protect multiple mammalian Na_V_ isoforms from STX block (Fig. 1) establishes that *Rc*Sxph possesses key favorable biophysical properties to counteract STX poisoning *in vivo*.

**Table 1:**
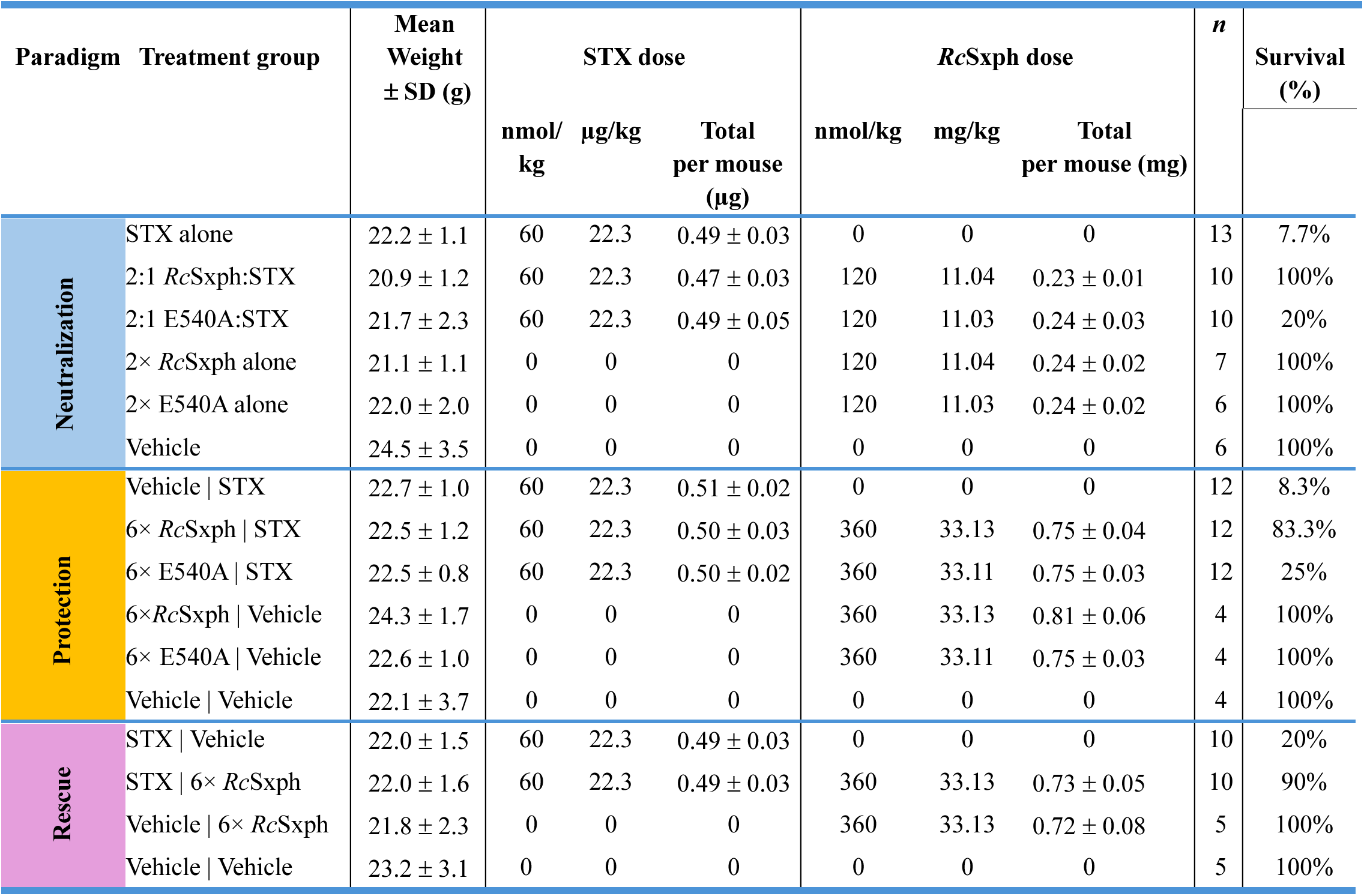
Summary statistics of treatment groups for STX neutralization, protection and rescue paradigms.

### Saxiphilin counteracts STX lethality in multiple administration paradigms

The presence of Sxph in frogs that are resistant to STX poisoning^6^, together with the ability of Sxphs to protect various Na_V_s from STX block (Fig. 1)^6,26,30^ has suggested that Sxph is a ‘toxin sponge’^6^ that acts *in vivo* to counter the lethal effects of STX. To test this idea directly, we established three paradigms in mice designed to investigate the ability of *Rc*Sxph to mitigate the neurotoxic effects of STX. We first investigated a ‘neutralization paradigm’ in which a 2:1 molar ratio of Sxph and STX were allowed to interact for at least 30 minutes prior to a single intraperitoneal (i.p.) injection of the equilibrated mixture (Fig. 2A). Control experiments injecting STX alone at 2.5× the reported LD_50_ (60 nmol/kg, i.p.)^34,35^ caused almost all mice (92.3%, total *n* = 13) to develop poisoning symptoms within three minutes with rapid progression to death soon thereafter. The median time of death (TOD) for mice treated with STX alone was 4.38 min (95% confidence interval (CI): 3.75–4.5 min, Fig. S3), aligning with the rapid lethality of similarly high STX doses reported in other studies^34–36^. Importantly, i.p. injection of *Rc*Sxph or E540A alone at 120 nmol/kg, corresponding to 2× the molar ratio of STX, caused no adverse effects and yielded survival outcomes (100% survival, total *n* for each group = 6) that were not different from vehicle controls (Figs. 2A and 2D, Table 1). Thus, both *Rc*Sxph and E540A appear well tolerated when delivered by this route and do not evoke undesirable reactions that could interfere with assessing their abilities to counteract the lethal effects of STX.

**Figure 2.**
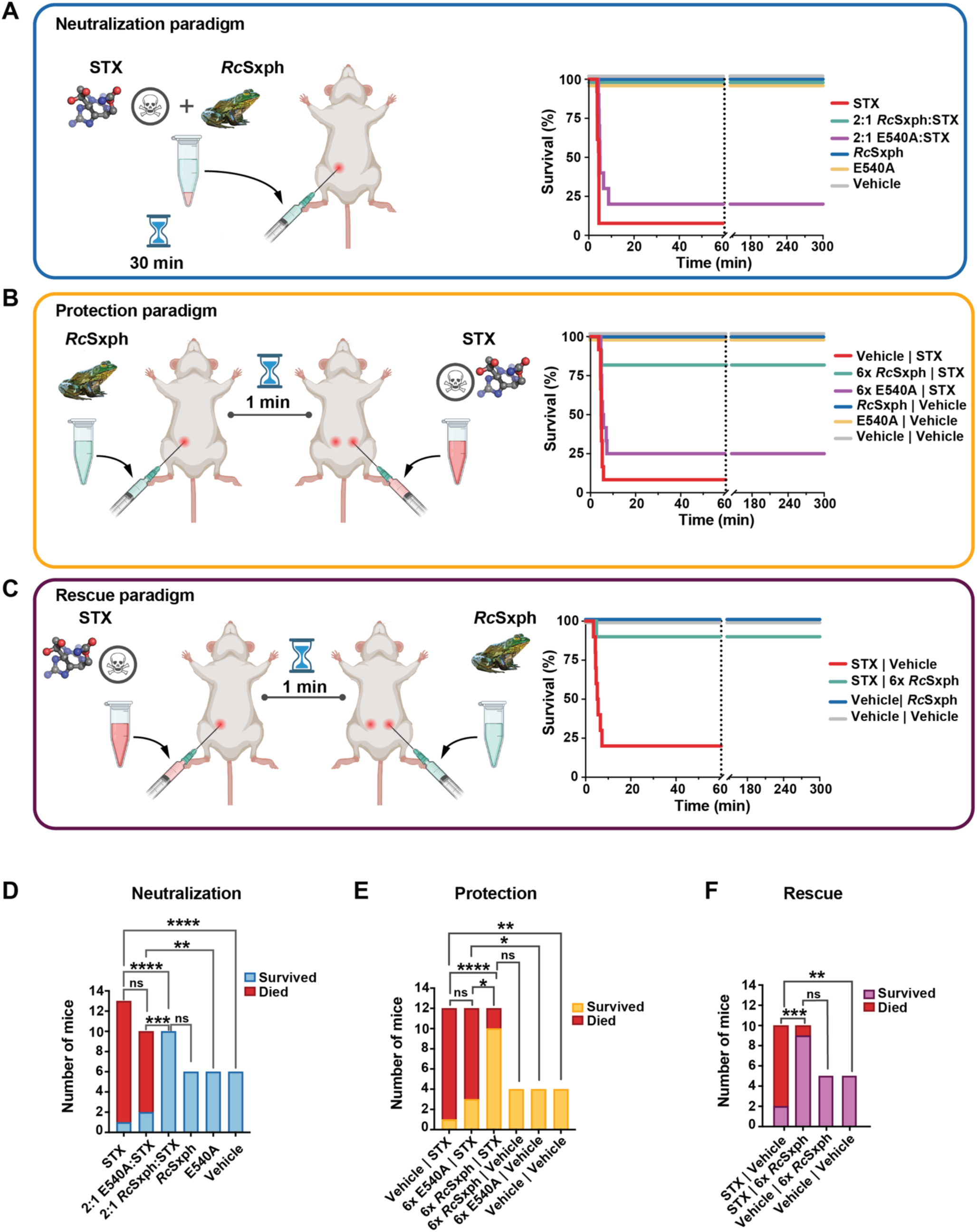
*Rc*Sxph abrogates lethality in a mouse model of severe STX poisoning under three different treatment scenarios. Schematics of experimental protocols created with Biorender.com and corresponding Kaplan-Meier survival graphs for: **A, Neutralization**, mice received a single i.p. injection of 2:1 [Sxph]:[STX], preincubated for 30 min; **B, Protection**, mice first received Sxph and then were challenged with STX 1 min later; and **C, Rescue**, mice first received STX and then Sxph 1 min later. All experimental compounds were administered via i.p. injection. Across all experiments, STX control was dosed as 2.5× LD_50_ (60 nmol/kg, 22.3 µg/kg) (red, dashed line). The maximum observation period for STX alone control mice was 60 min, indicated with a grey, dashed vertical line. Mice which received STX and *Rc*Sxph (green, solid) and STX and E540A (purple, dashed) were permitted to be observed for maximum 5 hours. All vehicle controls (grey, solid), and Sxph only controls (*Rc*Sxph only: blue, dashed; E540A only: yellow, solid) survived for the duration of the experiment (5 hours) and are overlapping at 100%. **D-F,** Mouse survival in **D,** Neutralization (blue), **E,** Protection (yellow), and **F,** Rescue (purple) paradigms. Fatality (red) is indicated for all three. Survival proportions were compared using Fischer’s exact test. The sample size for each treatment group in each scenario is reported in the Materials and Methods and Table S4.

Remarkably, all mice injected with a pre-equilibrated 2:1 *Rc*Sxph:STX mixture in the neutralization paradigm survived for the entire 5-hour observation period of the experiment (100%, *n* = 10) (Fig. 2A, Table 1, Supplementary Video S1), indicating that *Rc*Sxph was able to counteract the lethal effects of STX completely. By contrast replacing *Rc*Sxph with E540A, a mutant having severely compromised affinity for STX^26^, resulted in death for almost all of the mice. The E540A treated mice showed no significant difference in survival proportion (80% fatality, total *n* = 10, *p* = 0.56, Fig. 2D, Table 1) or TOD relative to STX controls (median TOD for 2:1 E540A:STX 4.91 min, 95% CI: 4.23 - 8.75 min, *p =* 0.1) (Figs. 2A and S3). Hence, unlike the outcome for *Rc*Sxph, mice receiving the E540A:STX mixture had no advantage over those receiving STX alone. Together, these neutralization paradigm results demonstrate that co-administration of *Rc*Sxph and STX is sufficient to protect against the lethal effects of STX. Moreover, as observed for *in vitro* rescue of Na_V_s (Fig. 1)^6,26,30^ this ability is absent from E540A and is entirely dependent on the presence of the high affinity STX binding site on Sxph.

To test a more challenging paradigm in which the antidote (*Rc*Sxph) and toxin (STX) are delivered separately, we next asked whether prophylactic administration of *Rc*Sxph would act as an effective STX neutralizer in an experiment we termed the ‘protection paradigm’. We delivered doses of *Rc*Sxph or E540A followed by STX as two contralateral i.p. injections separated by a 1-minute interval (Fig. 2B). For these experiments we used the same 2.5× LD_50_ STX dose as in the neutralization studies but increased the amount of Sxph to a molar ratio of 6× relative to STX for both *Rc*Sxph (denoted ‘6× *Rc*Sxph’) and E540A (denoted ‘6× E540A’) (360:60 nmol/kg, Sxph:STX i.p.) to compensate for possible differences in absorption and distribution rates between the small molecule toxin (∼300 Da) and ∼300-fold larger macromolecule (∼91 kDa). Control experiments in which the vehicle solution was injected first followed by 2.5× LD_50_ STX resulted in fatality for nearly all subjects (91.7%, total *n* = 12) (Figs. 2B and 2E) that matched the results from a single injection of STX alone (Fig. 2D, Table 1). These results indicate that the contralateral injection of fluid into the peritoneal cavity before toxin delivery did not affect the toxin lethality (median TOD for Vehicle | STX 4.77 min, 95% CI: 4.47–5.26 min, *p =* 0.1, Table S3). Investigation of the Sxph only controls showed that the 6× *Rc*Sxph and 6× E540A doses were well tolerated. Neither form of Sxph showed any apparent acute toxicity or other differences from vehicle injected controls (*n* = 4 each for 6× *Rc*Sxph and 6× E540A alone, Figs. 2B and 2D, Table S4), analogous to outcomes for the 2× Sxph doses administered in the neutralization paradigm. Hence, the protection paradigm provides a clean background for assessing Sxph activity.

Strikingly, we found that prophylactic administration of 6× *Rc*Sxph robustly protected mice against STX, resulting in most of the treated mice (83.3%, total *n* = 12) surviving for the duration of the experiment (Fig. 2B and E, Table 1, Supplementary Video S2). By contrast, prophylactic administration of 6× E540A was ineffective at counteracting the lethal effects of the subsequent STX injection. Mice treated with 6× E540A prior to the STX challenge showed no significant difference in survival (75% fatality, total *n* = 12, *p* = 0.59) (Fig. 2B and E, Table 1) or median TOD (Figs. 2B and S3) relative to control mice in which vehicle alone was injected prior to STX (median TOD for 6× E540A | STX 4.95 min, 95% CI: 4.68 - 7.0 min, *n* = 9, *p* >0.99, Table S3). As with the neutralization paradigm, the inability of E540A to counter STX lethality validates that a competent high affinity STX binding site on *Rc*Sxph is necessary for this protein to function as an effective toxin sponge. Together, these data demonstrate that prophylactic i.p. administration of Sxph is an effective way to counter the lethal effects of STX.

Encouraged by the capacity of *Rc*Sxph to prevent STX lethality in both the neutralization and protection paradigms, we next investigated a third poisoning scenario that we termed the ‘rescue paradigm’ in which the order of administration of Sxph and STX was reversed from the protection paradigm (Fig. 2C). For these experiments, the 2.5× LD_50_ STX dose was injected i.p. first followed by contralateral i.p. injection of 6× *Rc*Sxph one minute later. As with other control experiments, most mice given the 2.5× LD_50_ STX dose died (80% fatality, total *n* = 10) (Fig. 2C and F) succumbing rapidly to the toxin (median TOD 4.9 min, 95% CI: 3.5–7.2 min, *n* = 8) (Fig. 2C-D, Fig. S3, Table 1). As observed in the protection paradigm, the dose of 6× *Rc*Sxph alone was well tolerated and had no adverse effect on survival (100% survival, *n* = 5) (Fig. 2C and F, Table 1). Most notably, the single i.p. injection of 6× *Rc*Sxph rescued almost all of the mice from the lethal effects of STX, with 90% of mice surviving to the experimental endpoint (total *n =* 10, *p* = 0.006) (Fig. 2C and F, Table 1, Supplementary Video S3). These findings show that *Rc*Sxph can counter the effects of STX even when the organism has been exposed to the toxin prior to antidote administration. Collectively, the outcomes from the neutralization, protection, and rescue test paradigms demonstrate that, as proposed^6,11^, Sxphs can act as ‘toxin sponges’ for STX. Moreover, the results show that that administration of Sxph alone is sufficient to negate the lethal effects of STX when either co-administered with the toxin or administered immediately prior or post toxin challenge in mice, demonstrating that Sxph is effective in both pre- and post-toxin exposure scenarios.

### Saxiphilin protects mice from STX poisoning symptoms

Given its efficacy against STX lethality, we next analyzed whether *Rc*Sxph delivery also affected the onset or severity of neurotoxic symptoms of STX poisoning across the three test paradigms. Analysis of video records of our experiments showed that mice treated with 2.5× LD_50_ STX developed a stereotyped progression of symptoms consistent with neurotoxic poisoning (Fig. 3, Table S4, Supplementary Videos S1-S3). These included: sudden behavioral arrest where the animal appeared to ‘freeze’ with motionless staring; limb paralysis with uncoordinated movement and hindlimb dragging and splaying; head and neck jerks; the animal lying prone on its belly accompanied by loss of muscle tone and balance; clonic and tonic-clonic seizures in which the mouse fell on its belly or side, often marked with severe limb convulsions and body tremors; wild jumping; barrel-rolls; abdominal breathing and gasping; bilateral exophthalmia; and tonic extension, leading to death. The severity of symptoms rapidly progressed from onset of mild symptoms (e.g. freezing, typically 2–3 min post injection), to moderate symptoms (e.g. hind limb paralysis and head and neck jerks, typically 2.5–3.5 min), to severe symptoms (e.g. convulsive seizures, typically 3–4.5 min) to death (typically, 3.5–5 min) (Fig. 3). Most of these symptoms can be seen in Supplementary Videos S1-S3. Symptom onset times were consistent across all three STX challenge paradigms (Fig. S4) and agree with previous descriptions of symptoms of high dose STX poisoning in mice^34,35^. We did not observe cyanosis but interpreted abdominal breathing and gasping as indicators of respiratory paralysis, the primary mechanism of STX lethality^21^. Notably, all mice that reached the abdominal breathing stage died. Importantly, mice that received control administration of *Rc*Sxph or E540A alone at either the 2× dose used in the neutralization paradigm or the 6× dose in the protection and rescue paradigms showed none of the above symptoms (Fig. 4) or any other adverse behavior, such as nocifensive licking or scratching and were indistinguishable from vehicle treated mice. Hence, the *Rc*Sxph proteins were well tolerated at all experimental doses and had no apparent adverse effect.

**Figure 3.**
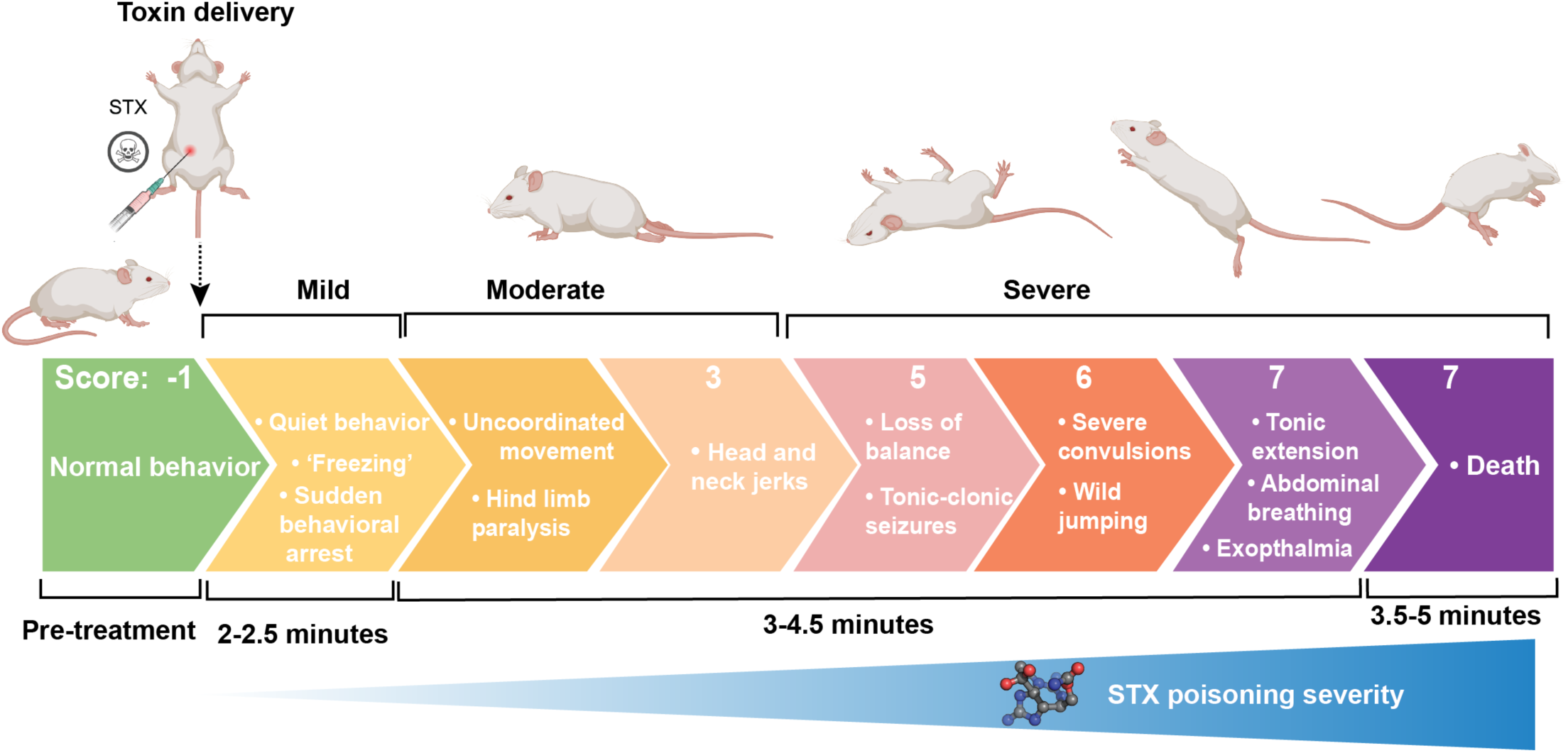
Symptom progression in STX poisoning. Schematic of typical symptom progression for STX poisoning for mice receiving 2.5× LD_50_ STX (60 nmol/kg) i.p.. Description of symptoms, typical onset times and corresponding revised Racine score of seizure severity ^37^ are indicated. Images created with Biorender.com.

**Figure 4.**
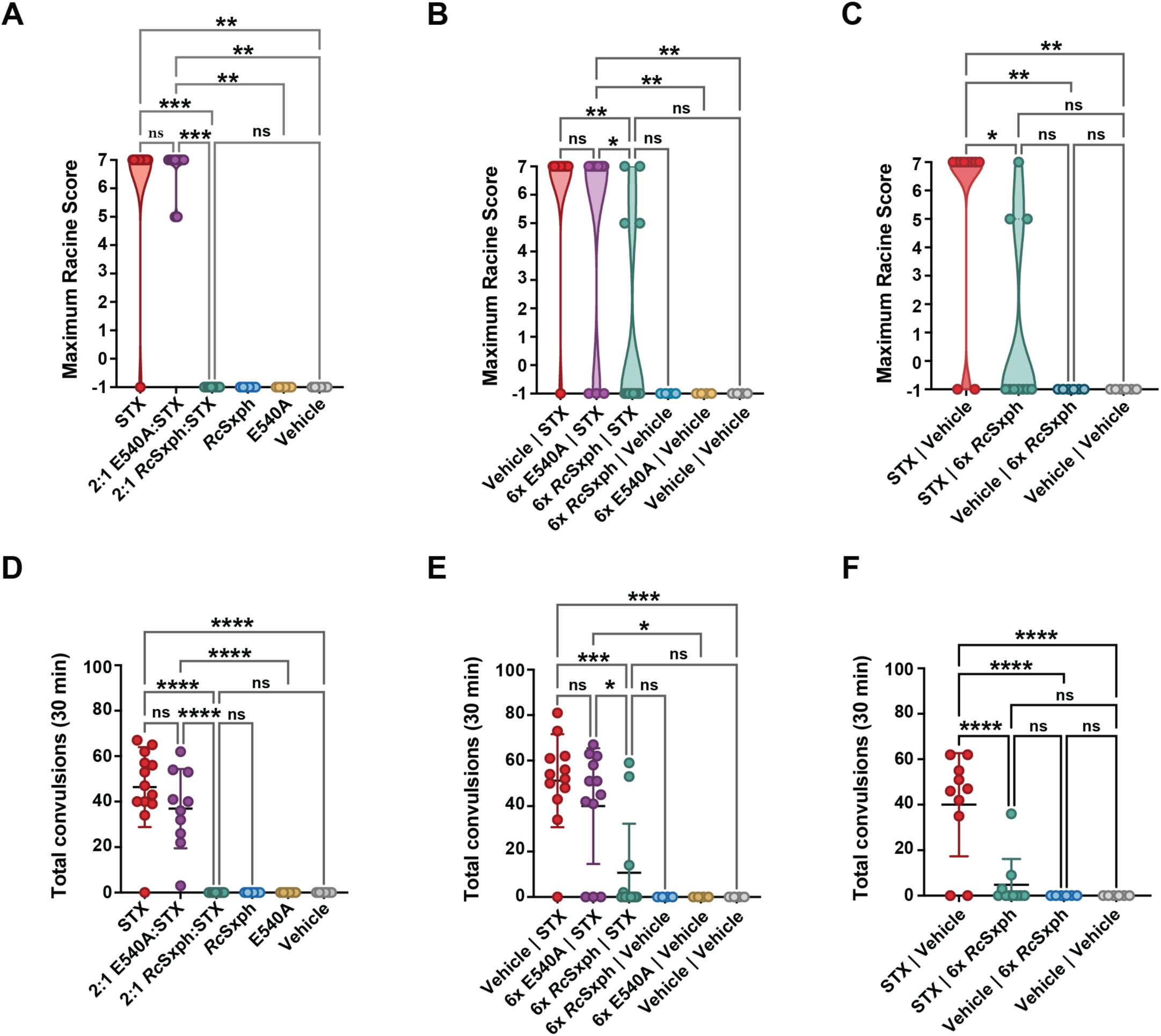
*Rc*Sxph ameliorates symptoms of severe STX neurotoxicity in mice. **A–C,** Mice seizures were scored using the revised Racine score described for PTZ-induced seizures in mice ^37^. Increasing score numbers reflect increasing seizure severity and are described in Table S3, with −1 representing normal behavior with no symptoms, and 7 reflecting fatal seizures. The maximum severity seizure score observed up to 30 min were plotted as violin-plots, with each point representing an individual animal. Asterisks indicate statistically significant differences between treatment groups (*p* < 0.05) using Kruskal-Wallis test with Dunn’s multiple corrections. **D–F**, Mouse convulsions (involuntary muscle contractions and jerks of the limbs, head or body corresponding to Racine score 3 or higher) were counted from video recordings in one-minute intervals for a maximum of 30 min following experimental compound administration. Each point represents the total convulsions counted for an individual animal, with the mean and standard deviation plotted. Asterisks indicate statistically significant differences between treatment groups (*p* < 0.05) calculated using one-way ANOVA with Tukey’s multiple comparisons test.

We found that the observed STX poisoning symptoms aligned with behavioral observations described for pentylenetetrazole (PTZ)-induced seizures^37^. This similarity enabled us to adopt the revised Racine scale for PTZ-induced seizures in mice^37^ (Table S3) to score the consequences of STX and Sxph treatment and quantitatively evaluate the outcomes of the different test paradigms. For this analysis, we focused on assigning seizure scores of 3 and higher, as these symptoms could be readily identified on video (Table S4, Supplementary Videos S1-S3). We tallied all involuntary jerks and spasms of the limbs, body, and head from video recordings to create a total convulsion count and used the maximum observed Racine score in combination with the total convulsions as markers of STX poisoning severity, and to assess Sxph intervention efficacy against these neurotoxic symptoms. This analysis showed that mice having seizures scoring ≥3 typically displayed progressively worsening convulsions that correlated with seizure duration and severity. All STX-treated mice that displayed abdominal breathing or developed seizures of score 6 progressed to death, regardless of treatment paradigm (Fig. 4A-C). In a few cases where Sxph was present, we identified mice that reached the clonic and tonic-clonic seizure stage (maximum score 5) but that ultimately recovered (Fig. 4A-C).

This quantitative analysis of video recordings from the neutralization paradigm (Supplementary Video S1) reinforced the observation that the 2:1 *Rc*Sxph:STX preincubation completely counteracted STX lethality (Fig. 2A). Further, mice receiving this mixture showed no indication of behavioral arrest, paralysis, seizures (Fig. 4A) or convulsions (Fig. 4D), indicating that the presence of *Rc*Sxph prevented development of neurotoxic symptoms associated with STX poisoning. By contrast, all mice given the preincubated 2:1 E540A:STX mixture developed poisoning symptoms (Fig. 4A and Supplementary Fig. S4A) and had numerous convulsions matching the STX control group (mean convulsions 36.9 ± 17.4 and 46.2 ± 18.3 respectively, *p* = 0.24) (Fig. 4D). Although most mice in the STX alone and 2:1 E540A:STX groups developed severe seizures that progressed to death (median Racine score 7 for both groups, *p* >0.999, Fig. 4A), one mouse from the STX only group showed no symptoms and no STX lethality. Two mice from the 2:1 E540A:STX group displayed paralysis and tonic-clonic seizures reaching a Racine score of 5, but slowly recovered within one hour post-toxin challenge (Figs. 4A and 4D, Supplementary Fig. S4A). The manifestation of severe poisoning symptoms in all 2:1 E540A:STX treated mice underscores the ineffectiveness of the Sxph having a compromised STX binding site to neutralize STX. By contrast, the remarkable effectiveness of *Rc*Sxph to counter poisoning symptoms in the 2:1 *Rc*Sxph:STX treatment group demonstrates the power of this ‘toxin sponge’ to tightly sequester STX and protect animals from its harmful effects.

Evaluation of the videos from the protection paradigm (Supplementary Video S2) revealed that prophylactic dosing of 6× *Rc*Sxph prevented development of STX poisoning symptoms in the majority of mice (66.6%, total *n =* 12, median Racine score = −1) (Figs. 4B and E, Supplementary Fig. S4B). There were two mice from the 6× *Rc*Sxph | STX group that had symptoms that progressed to death (Racine score 7, Figs. 2B and E, and 4B). Two other mice from this test group developed serious seizures, reaching maximum Racine score 5 (Fig. 4B) with the animals lying prone and having a few intermittent convulsions (Fig. 4E). Both then recovered within 15 minutes of reaching this stage and had no further symptoms for the remainder of the experiment (Supplementary Fig. S4). These outcomes were in stark contrast to both STX control mice and mice pre-treated with 6× E540A. In both groups, most mice developed STX poisoning symptoms (91.6% and 75% respectively, total *n* = 12 for both) and all mice that developed seizures progressed to death (median Racine score for both = 7, Fig. 4B). Accordingly, the maximum seizure severity and number of convulsions observed in mice receiving the prophylactic 6× *Rc*Sxph (median Racine score = −1, mean convulsions = 10.7 ± 21.6)(Fig. 4E) were significantly lower than those for the 6× E540A (mean = 40 ± 25.5, *p =* 0.01) and STX controls (mean = 51.2 ± 20.5, *p =* 0.0002). Notably, there was no difference between the 6× E540A group and STX controls in symptom frequency (*p =* 0.59), seizure severity (*p =* 0.41), or total convulsions (*p* = 0.75), further affirming that the high-affinity STX binding pocket is critical for preventing neurotoxicity and highlighting the efficacy of *Rc*Sxph as a neuroprotective agent against STX.

Reversing the administration order of *Rc*Sxph and STX in the rescue paradigm yielded equivalent levels of neuroprotection against STX symptoms that matched the protection paradigm (Figs, 4B-C, E-F, and Supplementary Figs. 4B-C). We found that 70% of mice treated with *Rc*Sxph remained asymptomatic and seizure and convulsion-free throughout the rescue experiment (Fig. 4C and F, Supplementary Fig. S4, total *n =* 10). Further, *Rc*Sxph intervention following STX delivery resulted in a significant reduction in seizure severity relative to STX control mice (median Racine scores of −1 and 7 respectively, *p =* 0.036, Fig. 4C) and a ten-fold reduction in total convulsions (mean = 4.8 ± 11.3 and 40 ± 22.7 respectively, *p* < 0.0001, Fig. 4F). In addition to one fatality, two mice receiving 6× *Rc*Sxph following STX administration developed serious seizures (Racine score = 5) with the animals laying prone but then recovered (Fig. 4C, Supplementary Fig. S4). The observation of instances of symptom progression and subsequent reversal in the protection and rescue paradigms suggests that *Rc*Sxph can reverse developing symptoms in individuals where there may be differences in the absorption of *Rc*Sxph relative to the toxin. Taken together, the quantitative analysis of symptom progression provides further evidence that *Rc*Sxph is a ‘toxin sponge’ whose action prevents STX poisoning and ameliorates the associated symptoms.

### Saxiphilin reaches different organs

The effectiveness of *Rc*Sxph, a ∼91 kDa soluble protein, as counteragent against a 300 Da small molecule toxin in our i.p. administration paradigms prompted us to investigate its biodistribution. To examine the fate of *Rc*Sxph, we used untargeted proteomics with label-free quantification (LFQ)^38^ to probe for the presence of *Rc*Sxph the kidney, heart, skeletal muscle, liver, and brain harvested at the experimental endpoint of five hours post-administration of an i.p. dose of *Rc*Sxph. We compared samples from mice treated with 6× *Rc*Sxph in the protection paradigm (*n* = 3–4) versus those receiving vehicle only (*n* = 2–3). *Rc*Sxph was robustly detected in all organs tested from mice dosed with 6× *Rc*Sxph with high sequence coverage spanning all Sxph domains (Table 2, Supplementary Fig. S5). The highest *Rc*Sxph abundance was found in intensely irrigated organs (rank order LFQ intensity kidney > heart ∼liver) (Fig. 5F, Table 2). Lower, but significant levels of Sxph were also detected in the brain (Fig. 5D and F) and in the hind limb skeletal muscle (Fig. 5E–F), a site where competition with the high affinity STX target Na_V_1.4. is expected to protect against the toxin-induced paralysis. Detection of *Rc*Sxph across varied tissues five hours after administration indicates that the protein persists *in vivo*, a property that consistent with its high thermal stability (Supplementary Fig. S2)^26,30^ and high number of disulfide bonds^26,27,30^. As expected, no Sxph signal was detected in any vehicle treated control tissues (Supplementary Table S5). The broad organ distribution indicates that *Rc*Sxph enters circulation disperses systemically following i.p. administration, consistent with absorption through the lymphatic system^39^. Interestingly, the observed distribution pattern mirrors the natural widespread preence of Sxph across varied frog organs^11,28,40–42^, suggesting that the ability to reach different organs is an inherent property of Sxphs. Comparing *Rc*Sxph and vehicle treated mice revealed minimal changes in overall protein abundance (Fig. 5A-E) with no proteins associated with acute stress-responses identified in the proteomes (Supplementary Table S5). This result is consistent with behavioral observations that *Rc*Sxph is well tolerated and does not promote adverse reactions within the five-hour window of the experiment. Together, these results demonstrate that i.p. administration of the anuran protein *Rc*Sxph leads to its wide systemic dispersion in a mammalian system. Such properties are likely important for its anti-STX effects.

**Figure 5.**
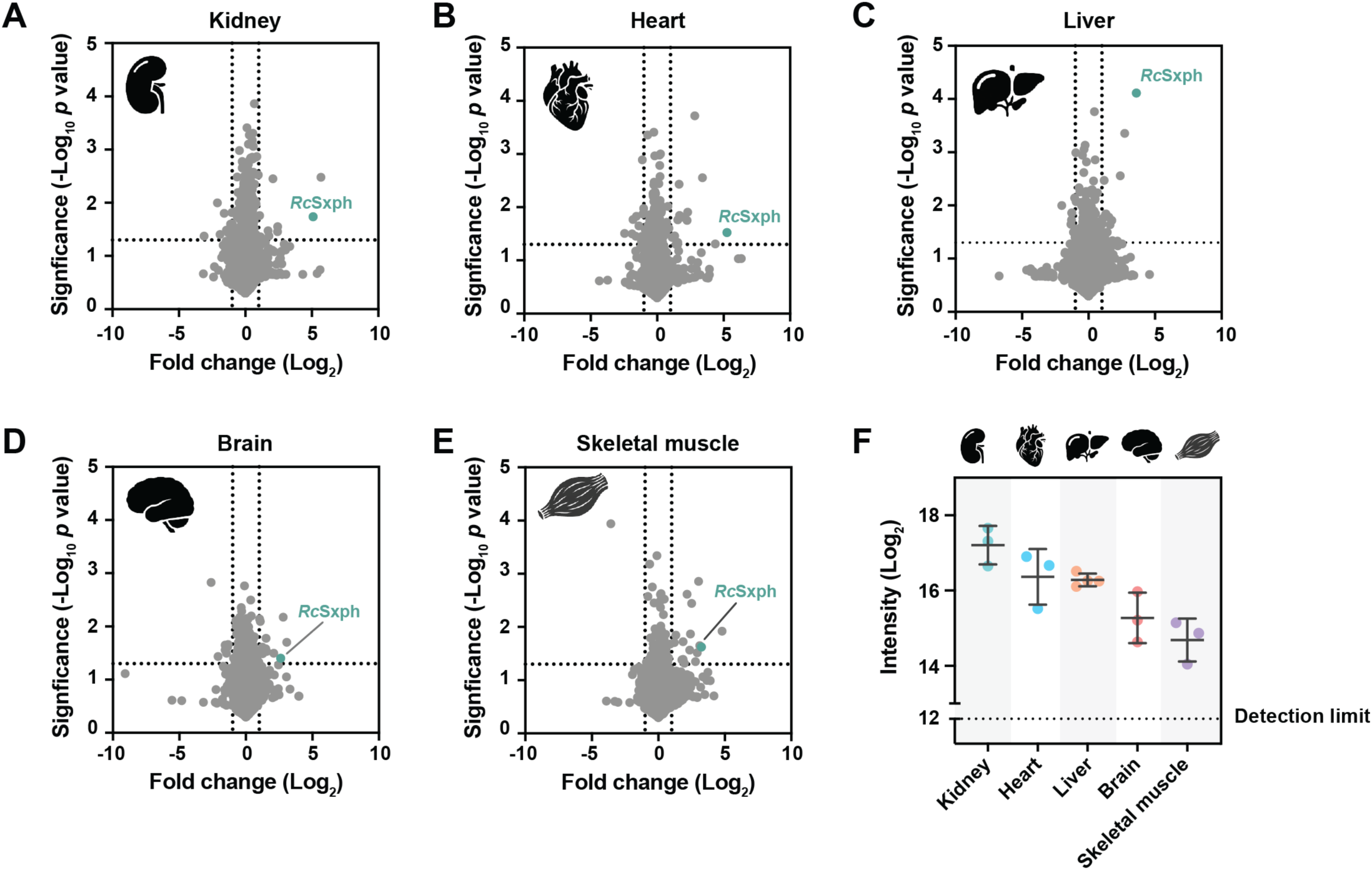
Mass spectrometry results of *Rc*Sxph treated mice 5 hours post-administration. **A-E** LFQ proteomic analyses of changes in protein expression from mouse organs harvested 5-hours post-i.p. administration of 6× *Rc*Sxph (360 nmol/kg) compared against vehicle only counterparts for **A,** kidney, **B**, heart, **C**, liver, **D**, brain and **E**, skeletal muscle. *Rc*Sxph is represented by green circles. Dotted lines indicate the thresholds for a 2-fold change in signal intensity (vertical) and statistical significance (*p* = 0.05, horizontal). **F** The intensity (log_2_) of *Rc*Sxph unique peptides for each tissue: kidney (green); heart (blue); liver (orange); brain (red); and skeletal muscle (blue). Each dot represents a biological replicate. Error bars represent SEM.

**Table 2:**
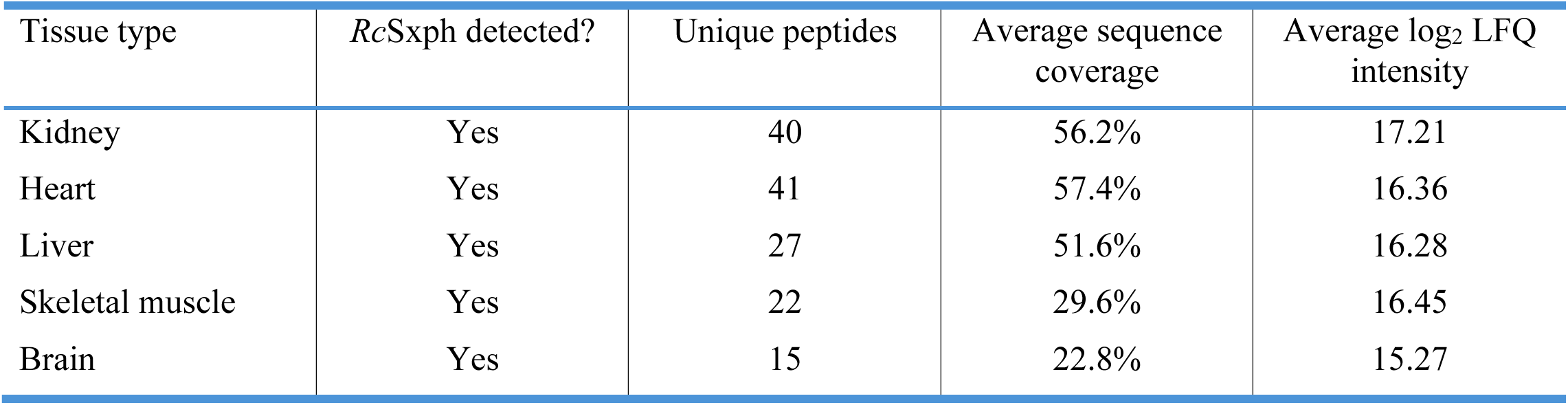
Summary statistics of LFQ proteomic analysis of mice treated with 6× *Rc*Sxph.

## Discussion

It has been known since the 1930s that frogs are resistant to STX poisoning^7^. This phenotype has been attributed to the presence of a family of soluble, high-affinity (Kd ∼nM) STX binding proteins called saxiphilins (Sxphs) that are thought to provide protection from STX poisoning by acting as ‘toxin sponges’^6,26,30^. Competition for toxin binding between ‘toxin sponges’ and their ion channel targets has been proposed as a generalizable toxin resistance mechanism^6,11,43^ that is distinct from classic ‘target site’ mutation strategies that rely on ion channel mutations^12–14^. Here, we investigated the ‘toxin sponge’ hypothesis using both *in vitro* assays and mouse models of STX poisoning. Consistent with previous electrophysiological studies examining the efficacy of different Sxphs against frog and human Na_V_1.4^6,26,30^, we found that *Rc*Sxph from the American bullfrog (*Rana catesbeiana*)^11,27^, the founding member of the saxiphilin family^26^, neutralizes the effects of STX against representative Na_V_s from human muscle, cardiac, and sensory nerves (*Hs*Na_V_1.4, *Hs*Na_V_1.5 and *Hs*Na_V_1.7) having varied STX affinities (Fig. 1). This broad protection for diverse STX targets supports the idea that Sxphs may be able to counteract STX exposure in animals that carry this protein. To investigate whether the robust *in vitro* ability of *Rc*Sxph to sequester STX would be sufficient to counteract the lethal effects of STX poisoning *in vivo*, we developed a set of three murine toxin challenge paradigms, denoted ‘neutralization’, ‘protection’, and ‘rescue’. Each paradigm involved i.p. injection in mice of 2.5× LD_50_ STX (60 nmol/kg), corresponding to ∼LD_90_, that induced the rapid onset of paralysis, seizures, convulsions, and death within 3–5 min (Figs. 2 and 3, Supplementary Fig. S4, Supplementary Videos S1-S3). Strikingly, a single dose of *Rc*Sxph negated both the neurotoxic and lethal effects of STX regardless of whether it was delivered together with STX (neutralization), before toxin administration (protection), or after toxin delivery (rescue). Such outcomes were not observed if *Rc*Sxph was replaced by the STX binding pocket mutant E540A that has a severely compromised STX binding capacity^26,30^. These findings confirm that Sxphs use this pocket to sequester the toxin *in vivo,* matching results from *in vitro* binding and functional studies^6,26,30^, and clearly establish that the ability of *Rc*Sxph to counter STX depends on its high affinity STX binding site. Together, these data show that Sxph alone is sufficient to prevent STX poisoning and validate the hypothesis that Sxphs are *in vivo* ‘toxin sponges’^6,11,26^.

For Sxphs to counteract STX poisoning the ‘toxin sponge’ must be able to sequester STX in a complex physiological setting. Accordingly, we observed that *Rc*Sxph was more effective against STX poisoning in the neutralization paradigm than in either the protection or rescue scenarios, likely reflecting disparities in when the toxin and *Rc*Sxph encounter each other versus the Na_V_ target. Administration of preincubated 2:1 *Rc*Sxph:STX prevented STX poisoning symptoms, such that all mice receiving this antidote:toxin mixture remained free from paralysis, seizures, and convulsions throughout the experiment and avoided death (Figs. 2A, 4A, and 4D, Supplementary Fig. S4A). The absence of neurotoxic symptoms indicates that the high-affinity Sxph:STX interaction established during preincubation was maintained to a sufficient level *in vivo* following i.p. injection to prevent minimal STX escape and interactions with Na_V_s. Strikingly, even though in the rescue experiment the antidote was delivered after toxin administration, both the protection and rescue paradigms revealed comparable levels of defense against neurotoxicity and death (67% asymptomatic, 83.3% survival; 70% asymptomatic, 90% survival, respectively). This excellent protection from the action of STX suggests that *Rc*Sxph is rapidly distributed in mice throughout the circulation following i.p. injection, a process that is likely to occur via absorption through the lymphatic system^39^. This interpretation is supported by mass spectrometry detection of *Rc*Sxph across multiple organs (Fig. 5, Table 2) that mirrors the widespread distribution of Sxph in frogs, where it is found not only in plasma^11,28^ but in varied organs^11,28,40–42^ including the kidney, liver, heart, muscle, and brain. Interestingly, despite its ∼91 kDa size, *Rc*Sxph was found in the brains of *Rc*Sxph-treated mice (Figs. 5D and F). Given the structural similarity between Sxphs and transferrins^27^, Sxph may be able to cross the blood-brain barrier via transferrin-receptor trafficking, an established route for macromolecules to enter the brain^44^. Establishing whether Sxphs interact with transferrin receptors or other cell surface molecules remains an important question. Importantly, our results indicate that *Rc*Sxph can access diverse tissues where it can capture circulating STX and consequently protect the animal from STX symptom development and lethality.

Because of its toxicity and designation as a chemical weapon^1,4^, there has been a long-standing interest in developing agents that can block the effects of STX^22,24^. However, classical antisera-based approaches to create agents that can bind STX or other guanidinium toxins have been challenged by the poor immunogenicity of this small molecule class^22,24^. Previous rabbit-derived STX antisera experiments in mice required either intravenous (i.v.) administration to rescue a lower-dose STX challenge^22^ or were only efficacious in preincubation neutralization experiments^24^. By comparison, here a single i.p. dose of *Rc*Sxph yielded higher survival rates against higher dose of STX than either of these prior efforts, with 100% survival in the neutralization paradigm and 90% survival when administered one-minute post-STX challenge (Figs. 2A and C). Hence, Sxph outperforms prior antisera-based approaches and highlights the potential of using Sxphs for STX countermeasure development. Recently, it was reported that *Rc*Sxph encapsulated in neuronal membrane nanosponges was efficacious against low dose subcutaneous (s.c.) injection of STX (4.5 µg/kg s.c.) in neutralization, protection and rescue scenarios^25^. Although these data support efficacy of Sxph against STX, it should be noted that the membrane encapsulation of these Sxph-loaded nanoparticles prevents Sxph exit^25^, making it unclear how the circulating toxin accesses the nanoparticle encapsulated Sxph to elicit the reported effects. Regardless, our results demonstrate that *Rc*Sxph alone effectively negates the lethality of a ∼5-fold higher dose of STX (22.3 µg/kg i.p.) across the three experimental paradigms without the need for encapsulation or i.v. administration. Importantly, we also show that *Rc*Sxph is effective against the development of STX neurotoxic poisoning symptoms (Fig. 4), a result that has not been reported in previous STX anti-toxin studies. Further, as a single biologic, *Rc*Sxph, or a derivative thereof, would be more amenable to cost-effective and scalable manufacturing than antisera approaches or neuronal membrane nanosponge formulations. Notably, *Rc*Sxph was well tolerated in mice without any apparent acute toxicity or adverse reactions throughout the observation period at doses up to 360 nmol/kg (∼700 µg in a 22 g mouse). This result together with the general high thermal stability of Sxphs (Supplementary Fig. S2, Table S2)^26,30^ underscores the potential of using this protein family for development of anti-PST agents.

Natural PST mixtures can contain >50 STX congeners, some of which are more toxic than STX^1,5,35,45^. Thus, in pursuit of effective PST antidotes, it will be important to advance toxin binding proteins that have a broad toxin binding capacity. In this regard, the STX-binding pocket in *Rc*Sxph and the closely related *Np*Sxph from *N. parkeri* have been shown to accommodate STX congeners having modifications such as decarbamoylation and mono- and disulfation while maintaining nanomolar binding affinities^30^. Similar to STX, this tight binding enables neutralization of toxin block in *in vitro* electrophysiology experiments^30^, suggesting that Sxphs may provide *in vivo* protection against poisoning by other STX congeners. The variability in the STX binding pockets of different Sxph family members^26^ as well as evidence that some animals may carry Sxph-like proteins capable of binding neo-STX congeners^31^ offers a platform for developing natural or designed proteins with enhanced toxin binding capacity and affinity. Further, the existence of other classes of toxin binding proteins in frogs^28,40,43^ and fish^46,47^ highlights the breadth of the ‘toxin sponge’ strategy as a biological mechanism of toxin resistance and raises the possibility this protein class can be leveraged as versatile source for developing antidotes for diverse classes of small molecule toxins. Together with recent advances in engineered binders for snake venoms^48^ and cholera toxin^49^, as well as development of anti-venom nanobodies^50^, our findings underscore the wide-ranging potential of anti-toxin binders as neutralizing agents against both protein and small molecule toxins.

STX and associated PSTs produced by dinoflagellates and cyanobacteria in marine and fresh water HABs are among the most potent neurotoxins known due to their strong interactions with Na_V_s that block bioelectrical signaling^1–3^. No PST antidote exists and management options for PSP events are limited to supportive care^8,9,21^. The increasing frequency, geographical range, and severity of HABs^15–17^ emphasizes the need to identify effective PSP antidotes. Our results demonstrate that it is possible to harness the American bullfrog *Rc*Sxph as an STX countermeasure and that administration of Sxph alone is sufficient to rescue and protect against the neurotoxic and lethal effects of STX poisoning, validating the idea that Sxphs act as ‘toxin sponges’ *in vivo*^6^. The diversity of the Sxph family^26,30^ provides a scaffold for development of variants having enhanced affinity and broad specificity for PSTs and points to the potential of capitalizing on the natural array of anti-toxin proteins^28,31,40,43,46,47^ as blueprints for developing toxin antidotes.

## Materials and Methods

### Saxitoxin preparation

STX used for whole-cell patch-clamp electrophysiology experiments was purchased as calibration-grade saxitoxin dihydrochloride from the National Research Council Canada (NRC, Halifax, Canada, cat. no. CRM-STX-g) and was lyophilized and redissolved in MilliQ water to make a 1 mM stock. STX used for animal studies was synthesized as ultra-pure saxitoxin dihydrochloride, purified, and validated as previously described^41^ and dissolved in MilliQ water to make a 5 mM stock. Toxin aliquots were frozen and stored at −20°C. Immediately prior to animal study use, STX stocks (200 µM) were freshly prepared in phosphate-buffered saline (PBS) (pH 7.4) (137 mM NaCl, 2.7 mM KCl, 10 mM Na_2_HPO_4_, pH 7.4) (bioWORLD, cat. no. 41620008).

### Saxiphilin expression and purification

*Rana catesbeiana* saxiphilin (*Rc*Sxph) (GenBank: U05246.1)^27^ and *Rc*Sxph E540A^26,51^ were produced as secreted proteins and purified as previously described^26,27,51^. Briefly, Sxphs carrying a C-terminal 3C protease cleavage site followed by green fluorescent protein (GFP) and a His_10_ tag and were expressed in *Spodoptera frugiperda* (*Sf9*) insect cells using a baculovirus expression system^51^. *Sf9* cells were infected with P2/P3 baculovirus at the dilution ratio of 1:50 (v/v) and incubated at 27°C in a non-humidified New Brunswick Innova 44 incubator (Eppendorf, cat. no. M1282-0010) with shaking at 130 rpm. After 72 h, cells were harvested by centrifugation (JLA-8.1000 rotor, 4000*g*). The supernatant was collected, adjusted to pH 8.0 with a final concentration of 50 mM Tris-HCl, and treated with 1 mM NiCl_2_ and 5 mM CaCl_2_ to precipitate contaminants. The resultant precipitants were removed by centrifugation (JLA-8.1000 rotor, 6200*g*). The clarified supernatant was incubated with antiGFP nanobody-conjugated Sepharose resin^52^ for 5 hours at room temperature (23 ± 2°C). The resin was washed with 20 column volumes of a wash buffer containing 300 mM NaCl and 30 mM Tris-HCl (pH 7.4). To remove the GFP-His_10_ tag, protein samples were treated on-column with 3C protease^52^ (0.2 mg mL^-1^ in the wash buffer) overnight at 4°C. The eluates were collected and purified by size exclusion chromatography (SEC) using a Superdex 200 10/300 GL column (Cytiva) in 150 mM NaCl,10 mM 4-(2-hydroxylethyl)-1-piperazineethanesolfinc acid (HEPES) (pH 7.4). Purified Sxph fractions were pooled and concentrated to 25-80 mg mL^-1^ using a 50-kDa cutoff Amicon Ultra centrifugal filter unit (Millipore, cat. no. UFC505096). Protein concentrations were determined by measuring UV absorbance at 280 nm using the extinction coefficient of 96,365 M^−1^ cm^−1^ for *Rc*Sxph and *Rc*Sxph E540A calculated using the ExPASY server (https://web.expasy.org/protparam/). Protein aliquots were flash-frozen in liquid N_2_ and stored at −80°C until further use. Prior to use in *in vitro* and *in vivo* studies, STX binding properties of each Sxph sample were validated using a fluorescence polarization binding assay^26,30,51^.

### *In vitro* toxin neutralization using patch-clamp electrophysiology

Planar whole-cell patch-clamp recordings were performed using a QPatch II platform (Sophion Bioscience). We used mammalian cells stably expressing the α-subunit of human Na_V_ isoforms: *Hs*Na_V_1.4 in HEK293 Griptite cells (SB Drug Discovery, SB-HEK-hNaV1.4); *Hs*Na_V_1.5, in HEK293 cells (SB Drug Discovery SB-HEK-hNaV1.5); and TE671 cells endogenously expressing *Hs*Na_V_1.7 (ATCC, RD-CCL-136)^53^. Cells were cultured according to manufacturer’s recommendations. The extracellular solution (EC, saline) contained: 2 mM CaCl_2_, 1 mM MgCl_2_, 10 mM HEPES, 4 mM KCl, 145 mM NaCl, 10 mM glucose (pH 7.4 with NaOH), and osmolarity adjusted to 305 mOsm/L with sucrose. The intracellular solution (IC) contained: 140 mM CsF, 1 mM/5 mM ethylene glycol-bis(β-aminoethyl ether)-N,N,N′,N′-tetraacetic acid (EGTA)/CsOH, 10 HEPES, 10 mM NaCl (pH 7.3 with 3M CsOH), and osmolarity adjusted to 320 mOsm/L with sucrose. Cells were dissociated using Detachin (AMSBIO, cat. no. T100100), washed and resuspended in EC to reach a cell density of ∼2.4 × 10^6^ cells/mL shortly before use. All experiments were conducted at room temperature (23 ± 2°C) using single-hole QPlates (Sophion Bioscience, cat. no. SB2040). All currents were digitized at 10 kHz and filtered at 3 kHz. Leak-subtraction protocol was applied, with non-leak-subtracted currents acquired in parallel.

Toxin concentration-response curves were determined by preparing 3-fold serial dilutions in EC and cumulatively applying increasing concentrations of toxin to the cells. A single pulse protocol was used with a 50 ms hyperpolarization step to −120 mV, followed by a 60 ms depolarization step from −120 mV to 0 mV, using a holding potential of –90 mV and a sweep-to-sweep interval duration of 5 s. The concentrations required to inhibit 50% of the current (IC_50_) were calculated by fitting the concentration-response curves (normalized peak current (I_x_/I_0_) as a function of STX concentration) using the following equation: I_x_/I_0_=(I_max_-I_min_)/(1+x/IC_50_), where I_x_ is the current amplitude at the STX concentration *x*, I_0_ is the current amplitude in the latest saline period before STX application, and I_max_ and I_min_ are the maximum and minimum peak current amplitudes, respectively, and IC_50_ is the half-maximal inhibitory concentration. We then calculated concentrations of STX sufficient to block ∼80% of the peak sodium current for use in subsequent *in vitro* neutralization experiments: *Hs*Na_V_1.4, 100 nM; *Hs*Na_V_1.5, 1.1 µM; and *Hs*Na_V_1.7, 3 µM.

For toxin neutralization experiments, stock solutions of *Rc*Sxph and *Rc*Sxph E540A (in SEC buffer, 150 mM NaCl and 10 mM HEPES, pH 7.4) were diluted in EC (0.3-9µM). Sxphs were incubated with STX using 3:1 [Sxph]:[STX] molar ratio for at least 30 min at room temperature (23 ± 2°C) prior application to the cells. To ensure that there was no voltage-dependent effect of toxin or Sxph, sodium currents were elicited using a double pulse protocol, with a holding potential of −90 mV, followed by a hyperpolarizing step to −120 mV for 50 ms, then a depolarizing step to 0 mV for 60 ms. Cells were then held at −70 mV for 200 ms and then depolarized to 0 mV again for 60 ms. As there were no significant voltage-dependent effects, the peak current from the first depolarization step was used for analysis. For each channel isoform, baseline sodium currents were first recorded in saline, and then test compounds applied in sequence: STX; Sxph:STX mixture; STX; and Sxph alone. The current recovery was then calculated as: I = (I_Sxph:STX_ – I_STX_)/(I_ctrl_ – I_STX_), where I_Sxph:STX_ is the current after application of Sxph:STX mixture, I_STX_ is the current after toxin application, and I_Ctrl_ is the baseline current recorded in saline. Statistical differences in current recovery were compared using unpaired t-tests with Welch correction. All data analyses were performed using the Sophion Analyzer Software (Sophion Bioscience) and GraphPad Prism v10.0 (GraphPad Software, Carlsbad, CA, USA).

### Isothermal titration calorimetry (ITC)

ITC measurements were performed at 35°C and 50°C using a MicroCal PEAQ-ITC calorimeter (Malvern Panalytical) as described previously^26,51^. *Rc*Sxph was purified using a final SEC step in 150 mM NaCl, 10 mM HEPES, pH 7.4. STX stock (1 mM) in MilliQ water was diluted with the SEC buffer to prepare 100 μM toxin solution having a final buffer composition of 135 mM NaCl, 9 mM HEPES, pH 7.4. To match buffers between the Sxph and toxin solutions, the purified protein samples were diluted to 10 µM with MilliQ water to reach a buffer concentration of 135 mM NaCl, 9 mM HEPES, pH 7.4. Protein samples were filtered through a 0.22 μm spin filter (Millipore, cat. no. UFC30GV00) before loading into the sample cell and titrated with 100 µM STX using a schedule of 0.4 μL titrant injection followed by 35 injections of 1 μL. The calorimetric experiment settings were: reference power, 5 μcal/s; spacing between injections, 150 s; stir speed 750 rpm; and feedback mode, high. Data analysis was performed using MicroCal PEAQ-ITC Analysis Software (Malvern Panalytical) using a fitted offset for correcting for the heat of dilution and the single binding site model.

### Animals and animal care

All animal experiments were approved by the Institutional Animal Care and Use Committee (IACUC), University of California, San Francisco (protocol approval #AN076215) and performed in accordance with University of California guidelines and regulations. Experiments were conducted using 5–6-week-old female CD-1 IGS mice (17–26.3 g, mean weight at dosing: 22.2 ± 1.7 g) sourced from Charles River Laboratories, USA. Mice were socially housed in groups of five, with environmental enrichment, food and water *ab libitum*, with 12/12-hour light cycles. Mice were habituated for at least 3 days prior to experiments with health monitored continuously. All mice used for experiments had body condition score 3^54^. Mice were allocated randomly to treatment groups.

### Mouse tests of Sxph efficacy against STX poisoning

Mice were weighed immediately prior to dosing. STX and Sxphs were prepared as nmol/kg doses in vehicle (PBS, pH 7.4) (bioWORLD, cat. no. 41620008) to a final volume of 100 µL and administered via intraperitoneal (i.p.) injection (using 28 gauge needle and 1 mL syringe). We used 2.5× the reported LD_50_ for STX (60 nmol/kg, 22.3 µg/kg) via i.p. injection in female mice (60 nmol/kg, 22.3 µg/kg)^34,35^. The Sxph dose was scaled (120-360 nmol/kg) according to the desired Sxph:STX molar ratio.

The primary outcome was survival. Survival proportions and time of death (TOD) were recorded for all mice. TOD was defined as the time at which the animal took its final gasping breath, per the Association of Official Analytical Chemists (AOAC) Paralytic shellfish toxin mouse bioassay (AOAC 959.08)^36^. Minimum sample sizes per treatment group were determined by using expected survival proportions in power analyses for comparing proportions of two independent populations. Values for significance (⍺) and power were set to 0.05 and 0.8 respectively. The survival proportions for STX only (*p*_1_) and Sxph intervention (*p*_2_) were conservatively approximated to 0.2 and 0.8, respectively, to give a minimum final sample size of *n* = 10. All power analyses were conducted using the University of British Columbia, Canada web-based calculator (https://www.stat.ubc.ca/~rollin/stats/ssize/b2.html)

Experiments were not blinded due to environmental health and safety and protocol requirements for STX handling. In accordance with IACUC protocol AN076215, mice that received STX only observed for maximum of 60 min post-injection, while mice that received either Sxph treatment or vehicle-only control were observed for up to five-hours post-injection. Animals that did not succumb to STX poisoning were euthanized via rising concentrations of CO_2_ followed by cervical dislocation at the defined protocol time endpoints.

### Sxph neutralization paradigm

For the neutralization paradigm in which Sxph and STX were pre-incubated prior to injection, Sxph:STX mixture doses were prepared in PBS to 100 µL as described above and incubated at room temperature (23 ± 2°C) for at least 30 min prior to administration as a single i.p. injection. Mice received either: (a) STX only (60 nmol/kg, *n* = 13); (b) 2:1 *Rc*Sxph:STX molar ratio (120 nmol/kg:60 nmol/kg, *n* = 10); (c) 2:1 E540A:STX molar ratio (120 nmol/kg:60 nmol/kg, *n* = 10); 2× *Rc*Sxph only (120 nmol/kg, *n* = 6); (d) 2× E540A only (120 nmol/kg, *n* = 6); or (e) vehicle only (*n* = 6). The mean body weight in this experiment was 21.6 ± 1.7 g.

### Sxph protection paradigm

For the protection paradigm, Sxph and STX were administered sequentially (Sxph first, STX second) as 100 µL contralateral i.p. injections separated by one minute. The Sxph:STX molar ratio for this experiment was 6:1 (360 nmol/kg:60 nmol/kg). Mice received either: (a) vehicle followed by STX (60 nmol/kg, *n* = 12); (b) 6× *Rc*Sxph (360 nmol/kg) followed by STX (60 nmol/kg, *n* = 12); (c) 6× E540A (360 nmol/kg) followed by STX (60 nmol/kg, *n* = 12); (d) 6× *Rc*Sxph (360 nmol/kg) followed by vehicle (*n* = 4); (e) 6× E540A (360 nmol/kg) followed by vehicle (*n* = 4); or (e) vehicle followed by vehicle (*n* = 4). The mean body weight of mice used in this experiment was 22.6 ± 1.4 g.

### Sxph rescue paradigm

For the rescue paradigm, STX and Sxph were administered sequentially (STX first, Sxph second) as 100 µL contralateral i.p. injections separated by one minute. The Sxph:STX molar ratio for this experiment was 6:1 (360 nmol/kg:60 nmol/kg). Mice received either: (a) STX (60 nmol/kg) followed by vehicle (*n* = 10); (b) STX (60 nmol/kg) followed by 6× Sxph (360 nmol/kg, *n* = 10); (c) Vehicle followed by 6× Sxph only (360 nmol/kg, *n* = 5); or (d) vehicle followed by vehicle (*n* = 5). The mean body weight of mice used in this experiment was 22.4 ± 1.9 g.

### Analysis of STX poisoning symptoms

In all experiments, mice were observed for normal baseline behavior for at least 30 min prior to dose administration, were observed for the full duration of the experiment in real time, and were video recorded for at least one hour post-dosing for further behavioral analyses. The following categories and time of onset of STX poisoning symptoms were recorded: (1) paralysis denoted by uncoordinated movement or hind limb dragging; (2) seizures of Racine score 3 or higher; (3) abdominal breathing; and (4) survival or death (TOD) defined as the time at which the animal took the final gasping breath^36^. Seizure severity was scored using the modified Racine score for PTZ-induced seizures in mice^37^ (Table S3), and the maximum score for each animal was recorded. We noted that animals that developed convulsions (sudden, irregular movements of the limbs or body due to involuntary muscle contractions, corresponding to seizures of Racine score 3 or higher) either succumbed to STX poisoning or stopped convulsing by ∼30 min. Therefore, total convulsions were counted by watching video recordings of mice in one-minute intervals for 30 min after the first experimental injection, and the total number of convulsions tallied and plotted. The supplementary video showing the STX poisoning symptoms was generated using iMovie v10.4.3.

### Statistical analysis of mouse rescue experiments

Efficacy of Sxph intervention in improving survival proportions in mice relative to STX controls was evaluated using Fischer’s exact test. Statistical differences in Kaplan-Meier survival curves between STX-control mice and *Rc*Sxph and E40A treated mice were compared using the Log-rank (Mantel-Cox) test. Differences between the medians of the maximum Racine scores for seizure severity and median TOD were compared using the Kruskal-Wallis test with Dunn’s multiple comparisons test. Statistical differences between treatment groups for total convulsions were determined using ordinary one-way Analysis of Variance (ANOVA) test with Tukey’s multiple comparisons test. All data were analyzed using GraphPad Prism v10.0 (GraphPad, Carlsbad, CA, USA).

### Proteomic analysis of saxiphilin distribution

Mouse organs selected for analysis were liver, kidney, heart, brain, and skeletal muscle, consisting of one entire gastrocnemius muscle. Mouse tissues were harvested by dissection immediately post-euthanasia. Excess blood was blotted and the organ washed with PBS. Whole tissues were then flash frozen with liquid nitrogen and stored at −80°C until use. Samples were shipped frozen on dry ice for subsequent proteomic analysis.

### Protein digestion

Tissue samples (liver, kidney, heart, brain and skeletal muscle isolated from the gastrocnemius muscle) were first washed with PBS, homogenized using a CryoMill cryogenic grinder (Retsch), sonicated using micro probe sonicator (Bandelin Sonopuls), and lysed by boiling at 95°C for 10 min in 100 mM Tris (pH 8.5) containing 2% sodium deoxycholate (SDC). Chloroacetamide and Tris(2-carboxyethyl)phosphine (TCEP) were added to 40 mM and 10 mM final concentrations, respectively. Protein concentration was determined using the Pierce bicinchonic acid (BCA) protein assay kit (Thermo Scientific). A final protein amount of 10 µg of per sample was used for the MS sample preparation. The sample volume was then adjusted to 65 µL in total by adding 100 mM triethylammonium bicarbonate (TEAB) containing 2% SDC.

Samples were further processed using SP3 beads on the KingFisher Flex automated Extraction & Purification System (Thermo Scientific) in a 96-well plate. Briefly, 65 µL of sample was added to 65 µL of 100% ethanol and mixed with the SP3 beads. After binding, the beads were washed three times with 80% ethanol. After washing, samples were digested in 50 mM TEAB at 40°C with 1 µg of trypsin for two hours, then another 1 µg of trypsin was added and digested overnight. After digestion, samples were acidified with trifluoroacetic acid (TFA, final concentration 1% (v/v)) and peptides were desalted using in-house made stage tips packed with C18 disks (Empore) as described^55^.

### nLC-MS/MS analysis

Nano-scale liquid chromatography-tandem mass spectrometry (nLC-MS/MS) was performed using nano reversed-phase columns (EASY-Spray column, 50 cm x 75 µm ID, PepMap C18, 2 µm particles, 100 Å pore size). Mobile phase A consisted of water and 0.1% (v/v) formic acid (FA). Mobile phase B consisted of acetonitrile (ACN) and 0.1% (v/v) FA. Samples were loaded onto the trap column (C18 PepMap100, 300 μm × 5 mm, 5 μm particle size, Thermo Scientific) for 4 min at 18 μL/min. The loading buffer was composed of water, 2% (v/v) ACN and 0.1% (v/v) TFA. Peptides were eluted with mobile phase B gradient from 4% to 35% B in 60 min. Eluting peptide cations were converted to gas-phase ions by electrospray ionization and analyzed on a Thermo Orbitrap Ascend (Thermo Scientific) by data independent approach. Survey scans of peptide precursors from 350 to 1400 m/z were performed in orbitrap at 60K resolution (at 200 m/z) with a 4 × 105 ion count target. DIA (data-independent acquisition) scans were performed in orbitrap at 30K resolution. AGC (automatic gain control) target was set to 1000% and maximum injection time mode to Auto. Precursor mass range 400 – 1000 m/z was covered by 30 windows 20 Da wide. Activation type was set to higher-energy collisional dissociation (HCD) with 28% collision energy

### Proteomic data analysis

All data were analyzed and quantified with the Spectronaut 18 software^56^ using directDIA analysis. Data were searched against a complete mouse proteome database (downloaded from uniprot.org in February 2024 containing 21,709 entries). Enzyme specificity was set as C-terminal to Arg and Lys, also allowing cleavage at proline bonds and a maximum of two missed cleavages. Carbamidomethylation of cysteines was set as fixed modification and N-terminal protein acetylation and methionine oxidation as variable modifications. FDR (false discovery rate) was set to 1% for PSM (peptide spectrum match), peptide and protein. Quantification was performed at the MS2 level. Precursor PEP (posterior error probability) cutoff and precursor and protein cutoff was set to 0.01, protein PEP was set to 0.05. Data were exported and data analysis was performed using Perseus 1.6.15.0^57^.

## Supporting information

Supplementary Figures S1-S5, Tables S1-S4, Descriptions of Table S5, and Videos S1-S3

## Data Availability

Mass spectroscopy data have been deposited at the Mass Spectrometry Interactive Virtual Environment (https://massive.ucsd.edu) under identifier MSV000099884. Data or materials will be provided on request from the corresponding author.

## Acknowledgements

The authors thank M. Huynh and M. Chagnon for technical support, D. Nagy and D. Sauter for technical advice, and K. Brejc for comments on the manuscript. LC-MS analyses were performed in the Laboratory of Mass Spectrometry at Biocev research center, Faculty of Science, Charles University, Czech Republic. This work was supported by United States Department of Defense grants HDTRA-1-19-1-0040, HDTRA-1-21-1-0011, and HDTRA-1-23-1-0026 to D.L.M., and NIH-NIGMS R01-GM117263-05 to J.D.

## Author Contributions

S.A.N., S.Z., S.J., Z.C., J.D., and D.L.M. conceived the study and designed the experiments. S.Z. and K.H. produced and purified Sxphs. S.A.N., S.Z., and A.B. established and performed electrophysiology experiments. Z.C. performed the ITC studies. S.A.N., S.Z., S.J., and K.H. conducted *in vivo* studies. D.R.G. and E.R.P. synthesized and quantified STX. S.A.N., S.Z., S.J., Z.C., and D.L.M. analyzed data. J.D., and D.L.M. provided guidance and support. S.A.N., S.Z., S.J., Z.C., J.D., and D.L.M. wrote the paper.

## Competing interests

J.D. is a cofounder and holds equity shares in TI Therapeutics, Inc., a start-up company interested in developing subtype-selective modulators of sodium channels.

## Notes

### Summary of Updates

Revision corrects errors in author affiliation and surname (Bara).

