## Supplementary Figures S1-S5, Tables S1-S4, Descriptions of Table S5, and Videos S1-S3 for "Saxiphilin functions as a ‘toxin sponge’ protein that counteracts the effects of saxitoxin poisoning"

4 December 2025

**Supplementary material for:**
**Saxiphilin functions as a ‘toxin sponge’ protein that counteracts the effects of saxitoxin poisoning**

Samantha A. Nixon[1#‡], Sandra Zakrzewska[1#], Seil Jang[1], Keli Huang[1], Anissa Bara[2], Zhou Chen[1†], Daynen R. Goss[3], Elizabeth R. Park[3], J. Du Bois[3], and Daniel L. Minor, Jr.[1, 4-8*]

[1]Cardiovascular Research Institute, University of California, San Francisco, CA
[2]Sophion Bioscience, Ballerup, Denmark
[4]Department of Biochemistry and Biophysics
[5]Department of Cellular and Molecular Pharmacology
[6]California Institute for Quantitative Biomedical Research
[7]Kavli Institute for Fundamental Neuroscience
University of California, San Francisco, CA 94158-9001 USA

[8]Molecular Biophysics and Integrated Bio-imaging Division
Lawrence Berkeley National Laboratory, Berkeley, CA 94720 USA

[3]Department of Chemistry
Stanford University, Stanford, CA 94305 USA

[#]Equal contribution

[‡] Present address:
Institute for Molecular Bioscience
The University of Queensland
St Lucia, QLD, Australia, 4067

[†]Present address:
Department of Anatomy and Physiology
Shanghai Jiao Tong University School of Medicine
Shanghai, 200025, China

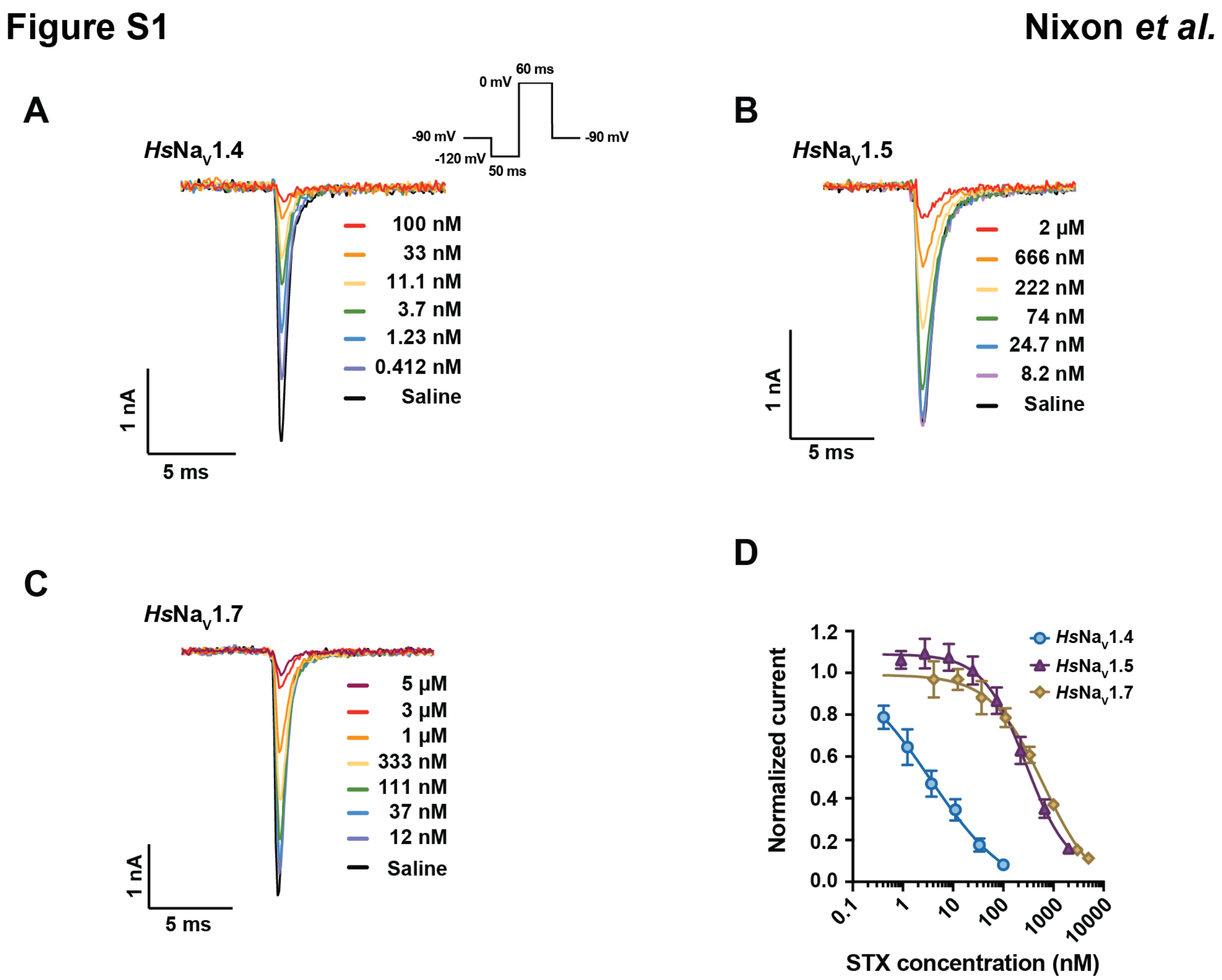


**Figure S1 Concentration-response curves for STX against human Navs. A-C,** Exemplar whole cell patch-clamp recordings of increasing concentrations of STX for **A,** *Hs*Nav1.4, **B,** *Hs*Nav1.5, and **C,** *Hs*Nav1.7. **D,** STX-response curves for *Hs*Nav1.4 (blue, circle, $n = 10$), *Hs*Nav1.5 (purple, triangle, $n = 9$), and *Hs*Nav1.7 (gold, diamond, $n = 6$). All points represent mean with SD.

**Figure S2** **Nixon *et al.***

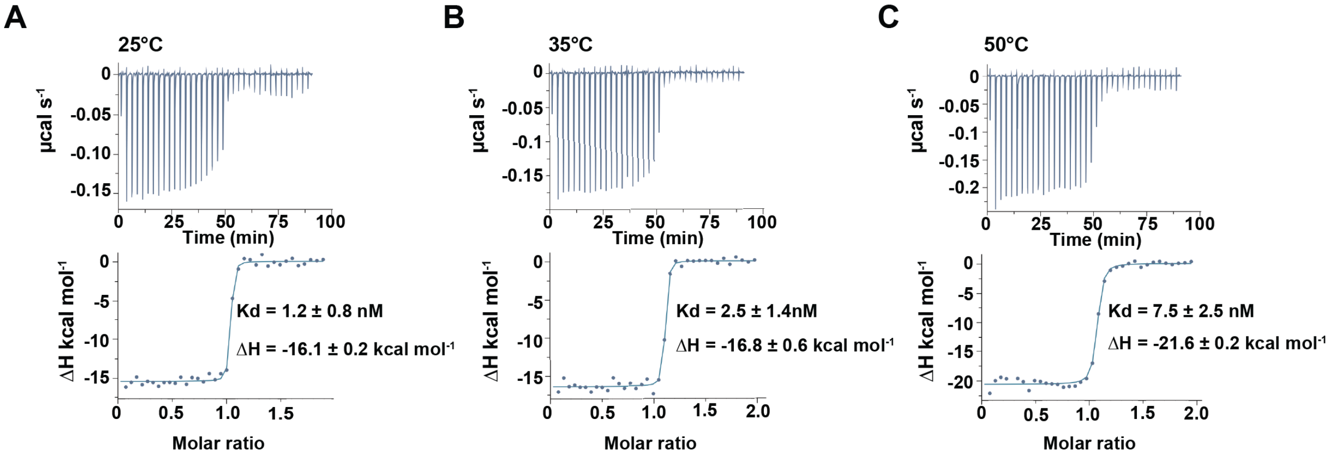


**Figure S2 ITC studies for *Rc*Sxph and STX at high temperatures. A-C,** Exemplar ITC isotherms for the titration of 100 μM STX into 10 μM *Rc*Sxph at **A,** 25°C (from [1]), **B,** 35°C, and **C,** 50°C**.** Kd and ΔH values are indicated.

**Figure S3** **Nixon *et al.***

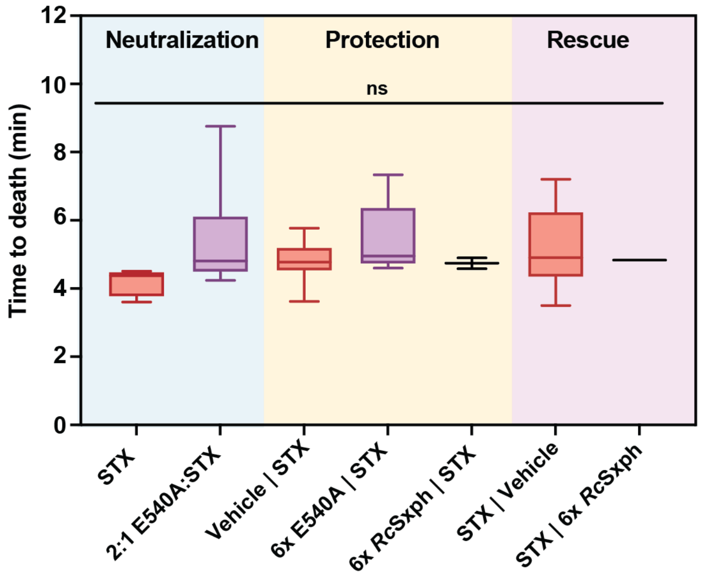


**Figure S3 Time to death for STX treated mice in neutralization, protection, and rescue paradigms.** Box plots showing the median, minimum, and maximum time to death (minutes) for each treatment group. The doses and number of fatalities in each treatment group were: Neutralization, STX alone (60 nmol/kg, $n = 12$), 2:1 E540A:STX (120 nmol/kg:60 nmol/kg, $n = 8$); Protection, Vehicle | STX (60 nmol/kg, $n = 11$); 6× E540A |STX (360 nmol/kg, 60 nmol/kg, $n = 9$); 6× *Rc*Sxph | STX (360 nmol/kg | 60 nmol/kg, $n = 2$); Rescue, STX|PBS (60 nmol/kg, $n = 8$), STX|6× *Rc*Sxph (60 nmol/kg | 360 nmol/kg, $n = 1$). Statistical differences between median time of death were compared using Kruskal-Wallis test with Dunn's multiple comparison test, no significant difference between groups was found ($p > 0.05$).

**Figure S4** **Nixon *et al.***

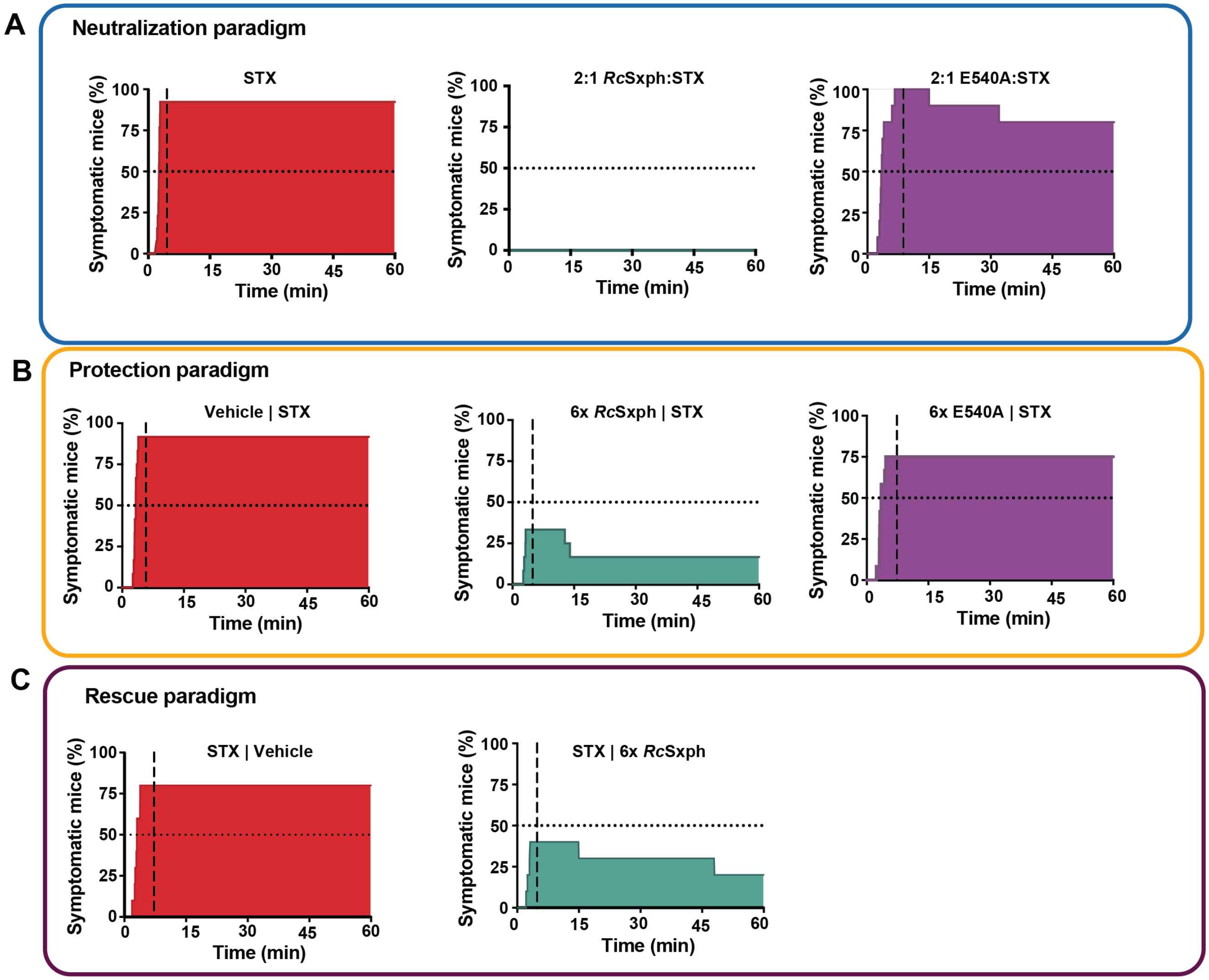


**Figure S4 Impact of Sxph treatment on STX poisoning symptoms and recovery over time.** Mice displaying STX poisoning symptoms, counted from initial symptom onset (min) were plotted as a percentage of the total treatment group for **A,** *neutralization*, **B,** *protection* and **C,** *rescue* scenarios. The vertical grey dashed line represents the median TOD for each group. Mice were counted as recovered when they had displayed no symptoms for at least 2 minutes and did not develop any further symptoms during the observation period. Mice that died were counted as symptomatic.

**Figure S5** **Nixon *et al.***

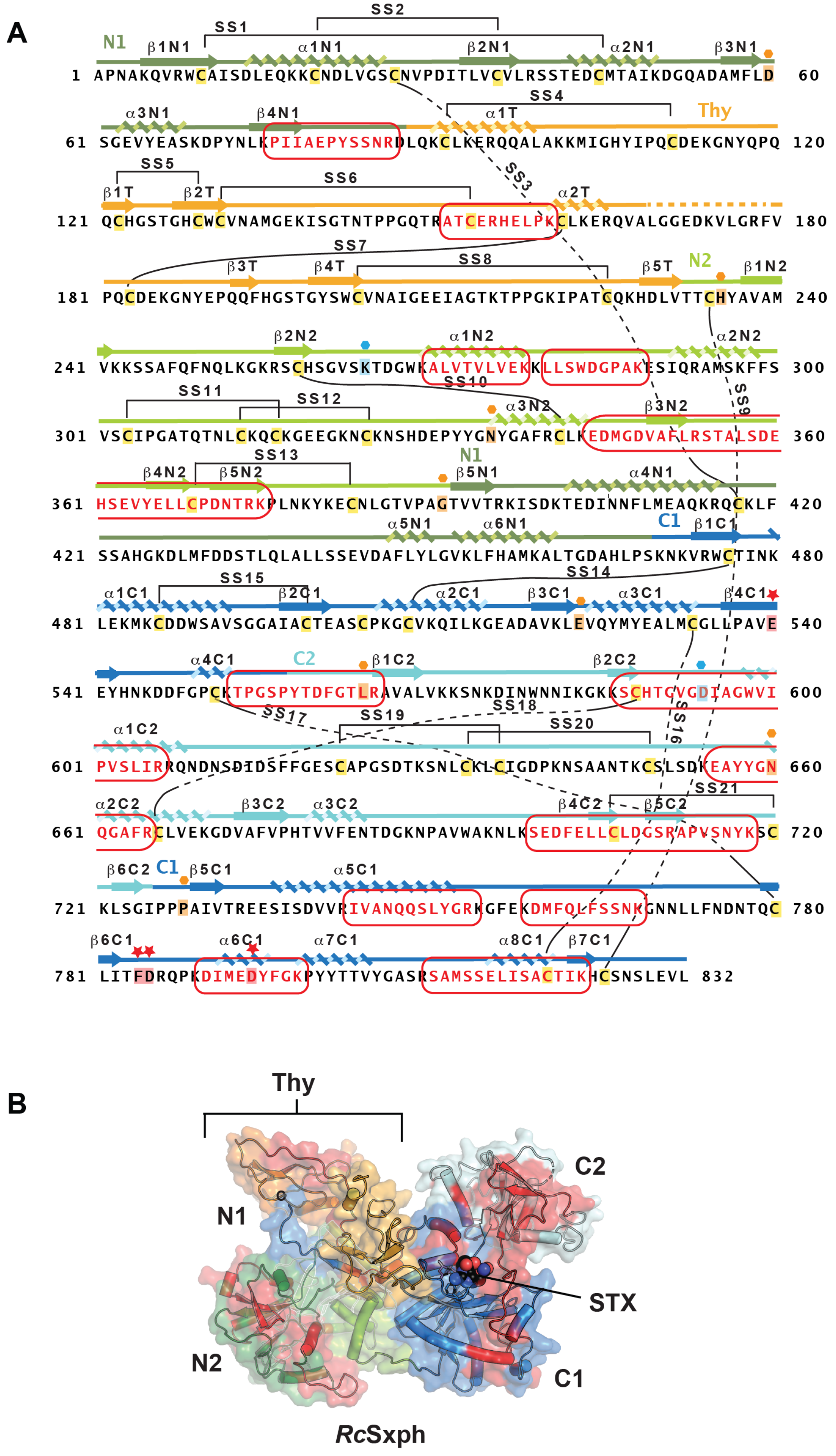

**Figure S5 Map of observed *Rc*Sxph peptide fragments A,** Shared observed peptides from mouse organs following 5-hour *Rc*Sxph treatment. **B,** Location of peptides from 'A' (red) mapped on the *Rc*Sxph:STX structure (PDB: 6O0F)[2]. Domains are indicated and are colored as follows: N1 (smudge), N2 (limon), thyroglobulin (Thy; bright orange), C1 (marine), and C2 (cyan). STX (black) is shown as space filling.

| **Table S1 STX $IC_{50}$s for *Hs*$Na_V$1.4, *Hs*$Na_V$1.5 and *Hs*$Na_V$1.7 and STX and Sxph concentrations used for *in vitro* neutralization studies** | | | | |
|---|---|---|---|---|
| **Isoform** | **$IC_{50}$ (nM)** | ***n*** | **[STX] [Sxph] *in vitro*** | |
| | | | **STX (nM)** | **Sxph (nM)** |
| ***Hs*$Na_V$1.4** | 3.5 ± 2.5 | 10 | 100 | 300 |
| ***Hs*$Na_V$1.5** | 284.6 ± 50.[illegible] | 9 | 1100 | 3300 |
| ***Hs*$Na_V$1.7** | 637.9 ± 189 | 6 | 3000 | 9000 |

$IC_{50}$, half-maximal inhibitory concentration

*n,* number of cells

Errors are SD.

**Table S2 *Rc*Sxph:STX thermodynamic binding parameters**

| Temperature | N (sites) | Kd (nM) | ΔH (kcal $mol^{-1}$) | ΔS (cal $mol^{-1}$ $K^{-1}$) | ΔG (kcal $mol^{-1}$) | ΔΔG (kcal $mol^{-1}$) | n |
|---|---|---|---|---|---|---|---|
| **25°C*** | 1.02 ± 0.01 | 1.2 ± 0.8 | -16.1 ± 0.2 | -12.7 ± 0.9 | -12.3 ± 0.5 | - | 3 |
| **35°C** | 1.02 ± 0.01 | 2.5 ± 1.4 | -16.8 ± 0.6 | -15.1 ± 3.3 | -12.2 ± 0.4 | 0.1 | 2 |
| **50°C** | 1.01 ± 0.06 | 7.5 ± 2.5 | -21.6 ± 0.2 | -29.7 ± 0.6 | -12.0 ± 0.0 | 0.3 | 2 |

N, number of binding sites

Kd, dissociation constant

$\Delta\Delta G = \Delta G_{T(°C)} - \Delta G_{25°C}$

n, number of observations

* Data taken from [1]

Errors are SD.

**Table S3 Median time of death (TOD) for STX treated mice in neutralization, protection, and rescue paradigms**

| Paradigm | Treatment group | Median TOD (min) | 95% CI | *n* |
|---|---|---|---|---|
| **Neutralization** | STX alone | 4.38 | 3.75–4.5 | 12 |
| | 2:1 E540A:STX | 4.8 | 4.23–8.75 | 8 |
| **Protection** | Vehicle \| STX | 4.77 | 4.47–5.26 | 11 |
| | 6x E540A \| STX | 4.95 | 4.68–7.0 | 9 |
| | 6x *Rc*Sxph \| STX | 4.74 | 4.58–4.9 | 2 |
| **Rescue** | STX \| Vehicle | 4.91 | 3.5–7.2 | 8 |
| | STX \| 6x *Rc*Sxph | 4.83 | nd | 1 |

nd = not determined due to insufficient *n*

95% CI, 95% confidence interval

*n* number of fatalities

**Table S4 Behavioral markers and associated seizure severity score**

| | Seizure Type | Score | Behavioral markers | Onset (min) | STX-treatment observational notes |
|---|---|---|---|---|---|
| Increasing seizure severity | **None** | -1 | Normal behavior | N/A | |
| | **Patrial/focal** | 0 | Whisker trembling | N/A | Whisker trembling was indistinguishable from control mice |
| | | 1 | Sudden behavioral arrest | 2–3 min | Observed in STX-treated mice. Not counted due to lack of correlative EEG. |
| | | 2 | Facial jerks | 2–3 min | Observed in STX-treated mice. Not counted due to lack of correlative EEG. |
| | **Generalized** | 3 | Head and neck jerks | 2.5–3.5 min | Observed in STX-treated mice. |
| | | 4 | Clonic seizure (sitting) | N/A | Not observed in any animals in this study |
| | | 5 | Clonic, tonic-clonic seizure with animal on belly (loss of muscle control) | 2.5–4 min | Observed. |
| | | 6 | Clonic, tonic-clonic seizure with animal lying on side or wild jumping | 3–4 min | Observed barrel-rolls in score 6 for STX-induced seizures |
| | **Death** | 7 | Tonic extension (laying down with limbs outstretched), death | 3–5 min | Tonic extension was accompanied by abdominal breathing, indicating respiratory paralysis |

Scores using modified Racine scale for PTZ-induced seizures in mice [3] with associated time of onset (min) following STX administration (60 nmol/kg i.p.) and observational notes.

Scores 3 and above were counted.

4 December 2025

**Table S5 Proteomic profiling of tissues from mice treated with 6× *Rc*Sxph or vehicle** Table lists all proteins identified across mouse kidney, liver, brain, heart and skeletal muscle by high-resolution mass spectrometry searched against a complete mouse proteome database (downloaded from uniprot.org in February 2024, containing 21,709 entries). The UniProt accessions, gene IDs, protein descriptions, q-value, C-score, and number of unique peptides are reported for each assigned protein. The LFQ intensity for each assigned protein is reported for each biological replicate. 6× *Rc*Sxph treated (shaded light green), Vehicle (unshaded).

**Supplementary Video S1 *Rc*Sxph neutralizes STX poisoning** Exemplar outcomes for neutralization of STX by *Rc*Sxph. Left mouse (green tail tag) was injected with a preincubated 2:1 molar ratio *Rc*Sxph:STX mixture (120 nmol/kg:60 nmol/kg i.p.). Right mouse (red tail tag) was administered with STX only (60 nmol/kg i.p.) as a control. The video shows symptoms of STX poisoning progressing to lethality in case of the STX-treated mouse. Timers indicate time post injection. The mouse injected with 2:1 *Rc*Sxph:STX survived and exhibits normal behavior 1-hour post-administration. Normal behavior continued for the remainder of the experimental protocol (maximum 5 hours, not shown).

**Supplementary Video S2 *Rc*Sxph prevents STX poisoning** Exemplar outcomes for protection from STX by *Rc*Sxph. Left mouse (green tail tag) was injected with 6× *Rc*Sxph (360 nmol/kg i.p.) followed by contralateral injection of STX (60 nmol/kg i.p.) after one minute. Right mouse (red tail tag) was administered a control solution of PBS (i.p.) followed by contralateral injection of STX (60 nmol/kg i.p.) after one minute. Timers indicate time following the second injection. The video shows symptoms of STX poisoning progressing to lethality in case of the STX-treated mouse. The mouse i.p. injected with 6× *Rc*Sxph | STX survived and exhibits normal behavior 1-hour post-administration. Normal behavior continued for the remainder of the experimental protocol (maximum 5 hours, not shown).

**Supplementary Video S3 *Rc*Sxph rescues STX poisoning** Exemplar outcomes for rescue from STX by *Rc*Sxph. Left mouse (red tail tag) was administered STX (60 nmol/kg i.p.) followed by contralateral injection of a control solution of PBS (i.p.) after one minute. Right mouse (green tail tag) was injected with STX (60 nmol/kg i.p.) followed by contralateral injection of 6× *Rc*Sxph (360 nmol/kg i.p.) after one minute. Timers indicate time following the second injection. The video shows symptoms of STX poisoning progressing to lethality in case of the STX-treated mouse. The mouse i.p. injected with STX | 6× *Rc*Sxph survived and exhibits normal behavior 1-hour post-administration. Normal behavior continued for the remainder of the experimental protocol (maximum 5 hours, not shown). Movie at 1 hour shows two mice from the STX | 6× *Rc*Sxph test condition.

4 December 2025
